# A diverse panel of 755 bread wheat accessions harbors untapped genetic diversity in landraces and reveals novel genetic regions conferring powdery mildew resistance

**DOI:** 10.1101/2023.03.23.533900

**Authors:** Rebecca Leber, Matthias Heuberger, Victoria Widrig, Esther Jung, Etienne Paux, Beat Keller, Javier Sánchez-Martín

## Abstract

Wheat breeding for disease resistance relies on the availability and use of diverse genetic resources. More than 800,000 wheat accessions are globally conserved in gene banks, but they are mostly uncharacterized for the presence of resistance genes and their potential for agriculture. Based on the selective reduction of previously assembled collections for allele mining for disease resistance, we assembled a trait-customized panel of 755 geographically diverse bread wheat accessions with a focus on landraces, called the LandracePLUS panel. Population structure analysis of this panel based on the TaBW35K SNP array revealed an increased genetic diversity compared to 632 landraces genotyped in an earlier study and 17 high-quality sequenced wheat accessions. The additional genetic diversity found here mostly originated from Turkish, Iranian and Pakistani landraces. We characterized the LandracePLUS panel for resistance to ten diverse isolates of the fungal pathogen powdery mildew. Performing genome-wide association studies and dividing the panel further by a targeted subsetting approach for accessions of distinct geographical origin, we detected several known and already cloned genes, including the *Pm2a* gene. In addition, we identified 22 putatively novel powdery mildew resistance loci that represent useful sources for resistance breeding and for research on the mildew-wheat pathosystem. Our study shows the value of assembling trait-customized collections and utilizing a diverse range of pathogen races to detect novel loci. It further highlights the importance of integrating landraces of different geographical origins into future diversity studies.

**Key Message:** A bread wheat panel reveals rich genetic diversity in Turkish, Pakistani, and Iranian landraces and novel resistance loci to diverse powdery mildew isolates via subsetting approaches in association studies.

## Introduction

Bread wheat (*Triticum aestivum* L.) provides more calories and protein per person than any other crop on Earth (FAO 2020). This allohexaploid (2*n* = 6*x* = 42, AABBDD) crop species originated from successive hybridization events, where the latest polyploidization is thought to have occurred ∼ 8,000 years ago in the Fertile Crescent (Glémin et al. 2019; Haas et al. 2019). After domestication, wheat was disseminated to Europe and Asia (Bonjean and Angus 2001), where accessions were selected based on the needs of individual farmers, thus becoming locally adapted traditional accessions, so-called landraces (Zeven 1998; Villa et al. 2005). With the Green Revolution in the 1960s, landraces were systematically replaced by advanced cultivars (Evenson and Gollin 2003) at the cost of a narrower genetic diversity due to the bottleneck effect of breeding (Tanksley and McCouch 1997; Reif et al. 2005). Consequently, modern cultivars are likely to lack a substantial proportion of the genetic diversity present in the wheat gene pool to combat abiotic and biotic stresses, including infection by pathogens. Plant diseases are a big threat to agriculture, with fungal pathogens playing a major role, causing an estimated annual yield loss of 18% in wheat (Savary et al. 2019). Among those, wheat powdery mildew, caused by the obligate biotrophic ascomycete *Blumeria graminis* f.sp. *tritici* (*Bgt*), is a major source of yield loss worldwide (Savary et al. 2019). Chemical control via pesticides is expensive and can negatively impact the surrounding ecosystem (Dormann et al. 2007; Bourguet and Guillemaud 2016). Besides, the European Commission proposed binding rules to reduce EU pesticide usage by 50% until 2030 (EU 2020). Therefore, alternative strategies are needed to control wheat mildew, especially in the context of climate change, as the geographical dispersal of pathogens and the severity of their infections are expected to increase rapidly (Singh et al. 2023).

In the Farm-to-fork strategy, the deployment of disease-resistant wheat accessions is proposed as a sustainable and effective way to combat pathogens (EU 2020). Such resistance can be conferred by resistance (*R*) genes, which typically encode intracellular nucleotide-binding leucine-rich repeat receptors (NLRs), albeit not always, that recognize pathogen avirulence effector proteins (AVRs) (Dodds and Rathjen 2010; Sánchez-Martín and Keller 2021; Athiyannan et al. 2022b). *R* gene resistance acts in a race-specific manner when an R protein recognizes the corresponding AVR (Flor 1971; Dodds and Rathjen 2010). The over 90 genetically characterized *R* genes against powdery mildew (*Pm* genes) represent a cornerstone of wheat breeding (McIntosh et al. 2013). However, race-specific resistance can be rapidly overcome by pathogens by evolving AVRs to evade recognition (McDonald and Linde 2002; Mundt 2014; Brown 2015). Due to this constant arms race and the low efficacy of the currently cloned *Pm* genes (Dracatos et al. 2023), the identification of new resistance genes is needed for wheat breeding programs. More than 7,000 distinct NLR-encoding genes are estimated to be present in the wheat gene pool (Walkowiak et al. 2020). Based on this genomic analysis, there are possibly hundreds of potentially active but unknown *Pm* genes in the wheat germplasm. The untapped genetic diversity of wheat landraces and their adaptation to individual environments with high disease pressure of locally adapted pathogens makes them promising candidates for containing such genes (Tanksley and McCouch 1997; Zeven 1998; Müller et al. 2018).

Together with cultivars and wild relatives, landraces are conserved in gene banks, where, to date, more than 800,000 *Triticum* accessions are stored (CGIAR 2023). However, these remain mostly uncharacterized for their potential in agriculture and their adaptive mechanisms are poorly understood, limiting their use in breeding (Tanksley and McCouch 1997; Müller et al. 2018). Attempts to unlock this hidden potential have been made using different approaches. For example, Balfourier et al. focused on maximizing the representation of different breeding statuses, dates of registration and geographical origin to reduce the collection size of the INRAe bread wheat collection from about 12,000 to 4,506 accessions (Balfourier et al. 2019). Genotyping these accessions improved understanding of wheat phylogeography and genetic diversity over time. Another study characterized the genetic diversity of 80,000 accessions, which represented a large part of the CIMMYT and ICARDA germplasm banks, covering not only domesticated hexaploid wheat but also tetraploids and crop wild relatives (Sansaloni et al. 2020). This revealed unexplored diversity in landraces and wheat selection footprints. In a third recent example, Schulthess et al. genotyped the IPK winter wheat collection of 7,651 accessions and a reference panel of 325 European elite cultivars (Schulthess et al. 2022). Later, this collection was phenotyped with a single powdery mildew isolate, detecting eleven previously undescribed resistance loci (Hinterberger et al. 2022).

Here, we used a panel based on former bread wheat collections assembled for allele mining (Bhullar et al. 2009, 2010b). These former collections included a main panel selected using a focused identification of germplasm strategy (FIGS) approach, revealing accessions with potentially high selection pressure for powdery mildew resistance (Mackay and Street 2004; Bhullar et al. 2009). We have now reduced these collections, focusing on landraces and maximizing the diversity of geographical origin. The reduced size of the panel allowed us to phenotype it with a diverse set of ten *Bgt* isolates, constituting a trait-customized panel that is ideal for searching for powdery mildew-resistant accessions and the underlying genes (Mascher et al. 2019). Using genome-wide association studies (GWAS) and a targeted subset approach for accessions of shared geographical origin and isolate-specific resistance patterns, we detected 22 most likely novel genetic regions associated with powdery mildew resistance.

## Materials and Methods

### Plant material and growth conditions

As starting material from which we selected our working panel, we utilized a formerly assembled bread wheat collection of 1,320 landraces that had been selected based on FIGS of accessions with potentially high selection pressure for powdery mildew resistance (Bhullar et al. 2009). Later, this collection was complemented with 733 accessions of diverse geographical origins (Bhullar et al. 2010b). Assessment of accession type, i.e., landrace, cultivar, breeding/research material or unknown, was based on passport data at https://www.genesys-pgr.org. When no GPS data was available for collection sites of accessions, we projected GPS using https://maps.google.com from the most detailed description available, i.e., given villages up to countries. Accessions from the two combined wheat collections were phenotyped for powdery mildew resistance at seedling stage with the six *Bgt* isolates CHE_94202, CHE_96224, CHE_97223, CHE_97266, CHE_98230 and GBR_JIW2. The infection phenotype was used to create a reduced panel of wheat accessions, consisting of approximately 50% that showed complete resistance (0% visible infection) to one or more isolates, or resistance with a threshold of 20% to at least two isolates. The remaining 50% were susceptible to all six isolates and had the same ratio of spring wheat to winter wheat as the resistant part of the panel. Additionally, we used wheat accession origin as a proxy for relatedness, choosing a geographically close susceptible counterpart for each resistant accession. The resulting diverse LandracePLUS panel of 755 bread wheat accessions (Table S1), with a focus on the fertile crescent, was infected with four additional *Bgt* isolates at seedling stage, i.e., CHE_19004, CHN_46_30, ISR_106 and ISR_94.

Differential lines used to assess virulence patterns for 27 different *Pm* genes are shown in Table S2, including near-isogenic lines (NILs) and accessions containing the designated gene. NILs had been backcrossed multiple times with susceptible accessions “Federation” or “Chancellor”, depicted by /x*Accession, where x is the number of backcrosses to the designated accession (McIntosh et al. 2013). Other differential lines were used as original seeds from the USDA ARS (https://npgsweb.ars-grin.gov/gringlobal/search) or propagated using isolation bags per single spikes.

Seeds used for infection tests were obtained by propagating accessions in the field using single rows per genotype without isolation. Seedlings for infection tests were grown in 40-well plastic trays in a growth chamber cycled at 20°C/16°C, 16/8h photoperiod with 80% relative humidity.

### Powdery mildew isolates and infections

We used previously sampled and sequenced *Bgt* isolates CHE_94202, CHE_96224, CHE_97223, CHE_97266, CHE_98230, CHN_46_30, GBR_JIW2, ISR_94 and ISR_106, which are described by Sotiropolous (Sotiropoulos et al. 2022). We sampled chasmothecia of one additional isolate, CHE_19004, in 2019 from a wheat field at Reckenholz, Affoltern, Switzerland, which was revived and sequenced as previously described (Sotiropoulos et al. 2022).

Powdery mildew infection tests of the differential lines and the LandracePLUS panel accessions were carried out on the primary leaves of ten to fifteen-day-old seedlings grown under the abovementioned conditions. Leaf segments were placed with their adaxial side up in Petri dishes filled with 0.5% Phyto agar containing 30 ppm benzimidazole. Fresh conidiospores were dispersed using 5 ml Pasteur glass pipettes in a settling tower (Lutz et al. 1992). Petri dishes with detached leaf segments were incubated for seven to nine days at 20°C, 80% relative humidity with a 16 h light/8 h dark cycle and 50 μmol m^-2^s^-1^ photon flux density. Infections were done in batches, with replicates of leaf segments from at least three independent seedlings per wheat accession on the same petri dish, infected at the same time. We used mildew susceptible accession Kanzler as a control for a proper mildew infection for all infection tests. Accessions Chancellor and Federation were used as additional susceptible controls for tests on differential lines. These susceptible controls were grown together with the tested accessions of each batch and distributed throughout the layout petri dish. If controls were not well infected and, in addition, the overall infection was low, the full infection test was repeated rather than using controls as a means to correct phenotypic values.

Disease levels were assessed seven to nine days after inoculation, depending on fungal growth in the batch, using a discrete quantitative scale with a score from 0 to 100 for the percentage of leaf area covered by sporulating mildew colonies, as described earlier by Kaur (Kaur et al. 2008). Disease levels of differential lines were directly scored as resistant (<=20) and susceptible (>20).

### DNA extraction and genotyping

DNA extraction of plant material was performed as previously described (Stein et al. 2001). DNA quality was assessed via agarose gels and genotyped using the TaBW35K single nucleotide polymorphism (SNP) array (Paux et al. 2022). The SNP call dataset included marker positions and flanking sequences based on RefSeq v1.0 of Chinese Spring (IWGSC 2018).

### *Pm* gene screening via polymerase chain reactions (PCRs) and sequencing

PCR analysis was performed using *Pm4* (Sánchez-Martín et al. 2021) and *Pm2* haplotype-specific markers (Manser et al. 2021). Four random landraces that were identified to carry *Pm2* were then used for long-range and a following nested PCR to amplify the gene for Sanger sequencing as previously described (Sánchez-Martín et al. 2016).

### General data analysis and visualization

Unless indicated otherwise, analyses were done using R version 3.6.3 (R Core Team 2022), including data handling with Tidyverse (Wickham et al. 2023) and visualizations with R package ggplot2 version 3.3.6 (Wickham 2016).

Kinship matrices for hierarchical clustering and visualization were done with GAPIT version 3 (Wang and Zhang 2021). Dendrogram formation and hierarchical clustering were performed using the stats R package version 3.6.3 (R Core Team 2022) functions hclust (method = ward.D2) and dist (method = euclidean). Defining clusters of genotypes was done using the package dendextend version 1.16.0 (Galili 2015).

Wheat *Pm* gene sequence assembly and alignment was done with CLC Genomics Workbench version 20.0.4 (Qiagen Bioinformatics, https://digitalinsights.qiagen.com/). Pathogen *Avr* gene sequence alignment was done using IGV version 2.15.4 (Robinson et al. 2011).

### SNP filtering and file format

We filtered for “PolyHighResolution” or off-target variants (OTVs) and markers with known chromosomal positions in the Chinese Spring RefSeq v1.0 reference genome. Thresholds of 25% heterozygosity, 25% missing data per wheat accession and 5% missing data per marker were applied, and markers with duplicated positions were removed. Absent haplotypes of an OTV were translated to “NA” to facilitate their inclusion in downstream analyses. Taken together, this resulted in 29,965 polymorphic markers. These were brought into a Hapmap format with R and then transformed into a variant call format (VCF) file using the software TASSEL version 5.0 (Bradbury et al. 2007). This file was used as an input to generate plink files using vcftools version 0.1.16 (Danecek et al. 2011), which were then transformed together with phenotyping data into .bed, .bim and .fam files using PLINK v1.07 (Purcell et al. 2007) for Admixture and association analyses.

### Phenotypic data analysis

The raw median of the biological replicates was taken as the final phenotype for seedling resistance assessment. Inconclusive phenotypes, e.g., 50% resistant and 50% susceptible against the same isolate due to possible seed contamination or heterozygosity, were excluded from further analyses. Phenotypes with less than three replicates were also excluded. These values were transformed into two categories, where 0-20% = resistant and >20% = susceptible for GWAS and Mantel tests. For Pearson correlation, phenotypic values of the differential lines were transformed to 0 and 1, respectively.

Pearson correlation between the isolate phenotypes was calculated using the stats package and visualized with the package corrplot version 0.92 (Wei and Simko 2021). Heritability was calculated for each pathogen isolate using R package lme4 version 1.1-34 (Bates et al. 2015) and a nested linear mixed model approach, where fixed variance is defined as the wheat genotype, and random variance as the infection test batch (“Round”) nested within the specific petri dish (“Plate”). P-values are based on ANOVA tests between the full and null models.

### *In silico* genotyping of TaBW35K array SNP variants in high-quality sequenced wheat genomes

For the comparison of genetic diversity, Fielder (Sato et al. 2021), Renan (Aury et al. 2022) and the 10+ wheat reference genomes ArinaLrFor, Chinese Spring, Claire, Cadenza, Jagger, Julius, Landmark, Lancer, Mace, Paragon, Norin61, Robigus, Stanley, SYMattis and Weebill (IWGSC et al. 2018; Walkowiak et al. 2020) were added. Flanking sequences of SNP array markers were queried using BLASTN searches against the publicly available wheat genomes. Blast results were filtered for hits with at least 60bp alignment and 96% shared identity, allowing no more than three mismatched nucleotides. Positions of the SNPs were then extracted from BLASTN alignments on the respective genomes through in-house scripts. Merging the resulting dataset with the 29,965 previously used markers resulted in an overlap of 27,337 SNPs used for principal component analysis (PCA) and hierarchical cluster analysis.

### Diversity Analysis via PCA and hierarchical clustering

PCAs for the LandracePLUS panel and high-quality sequenced genomes were done based on the 27,337 SNP set using the R package SNPRelate version 1.18.1 and gdsfmt version 1.20.0 (Zheng et al. 2012). For the PCA for the comparison to the 632 landraces from the INRAe study (Balfourier et al. 2019), we first filtered the genotyping data of the combined datasets provided by INRAe for the same cleaned 29,965 SNPs of the LandracePLUS panel. Because no creation of a Hapmap or VCF file was necessary for further downstream analysis, this dataset could be directly used for PCA using the prcomp function.

Hierarchical clustering analysis was performed using the kinship matrix of the 27,337 SNP set, including high-quality sequenced genomes. We visualized the dendrogram using dendextend version 1.16.0 (Galili 2015).

### Admixture kinship analysis and comparative visualization

We used the .bed file of 29,965 SNPs (not including the chromosome-assembled genomes) as an input to assess population structure using ADMIXTURE version 1.3.0 (Alexander et al. 2009), where the most likely number of founder populations K can be estimated via running the model over a series of values of K and then choosing K around the lowest occurring cross-validation (CV) error. We ran the model for K = 2 to K = 20. However, the CV error steadily dropped with increasing K (Fig S1). We, therefore, regarded Ks after the largest CV error drops of 33% in total as appropriate estimations for population structure, i.e., K = 4 to K = 6. A bootstrap of 500 and CV of 10 was used for this analysis.

A kinship matrix for the visualization was made with the same 29,965 SNP set using GAPIT version 3 (Wang and Zhang 2021). The kinship matrix was used as an input for hierarchical clustering for the comparative dendrogram. This dendrogram was visualized using the R package ggdendro version 0.1.23 (de Vries and Ripley 2022). Finally, these two plots and the admixture barplot were merged using the R package patchwork version 1.1.1 (Pedersen 2020).

### Mantel test

To calculate the Mantel test, we first transformed the VCF file of 29,965 SNPs into genlight format using the R package vcfR version 1.14.0 (Knaus and Grünwald 2017). Then, we used this as an input for producing a Bray-Curtis genetic distance matrix with the vegdist function from R package vegan 2.6-4 (Oksanen et al. 2022). This genetic matrix was then correlated to the Euclidean phenotypic distance matrices using the vegan package mantel function with the Spearman method and 999 permutations to obtain Mantel r values.

### Genome-wide association analysis (GWAS)

For GWAS, missing data from the 29,965 SNP set was imputed using general Beagle version 5.4 (Browning et al. 2018). GWAS and estimates of effect size – beta – were calculated using the GEMMA (Zhou and Stephens 2012) univariate linear mixed model. For the full LandracePLUS panel and each subset, .bed, .bim and .fam files were created as described above. .fam files were used as input for creating the kinship matrix in GEMMA using the options -gk 1 and -miss 1 . MAF was set to 1% for the full LandracePLUS panel and all subsets above 300 accessions in size, while for the other runs the MAF was set to 5%. This matrix was then integrated into the univariate linear mixed model with option -miss 1 (Zhou and Stephens 2012). Phenotypic data for the association studies was added to the .fam in R. P-values were based on the likelihood ratio test, and -log10 transformed for Manhattan plots. We used two thresholds to account for multiple testing: the False Discovery Rate (FDR) and the more conservative Bonferroni correction (BC). However, we regarded SNPs that passed the FDR test as significant.

Due to the design of the TaBW35K SNP array based on linkage disequilibrium (LD), meaningful LD decay analysis was not possible. We, therefore, decided to define GWAS peak intervals based on the LD decay results of a recent study on bread wheat using over 40 million SNPs (Liu et al. 2023). There, LD decay was found to be 6.0353, 2.3851 and 3.0278 Mb for subgenomes A, B and D, respectively. Hence, we defined peaks in a subgenome-dependent manner, as regions where at least two SNPs were significantly associated with the powdery mildew phenotype within the range of the above LD decay bp distances. We further considered significant single SNP associations as peaks if they uniquely mapped to one chromosome (weak homology on homoeologs, i.e., at least six additional SNPs/gaps), were surrounded by no or few SNPs and (1) they occurred for more than one isolate or (2) their significance passed the more stringent BC threshold. The most significant SNP – the peak SNP – plus/minus the subgenome-specific LD decay distance was used to define each peak interval. Alternatively, if several peak SNPs occurred, both were used for the interval calculation.

To test if the peak on chromosome 5D was derived from *Pm2* and account for its presence, we included a covariate (option –c) for binary information on the presence of *Pm2* based on haplotype-specific PCR screening.

Random subsets were produced using R package tibble 3.1.8 (Müller and Wickham 2022). Subsets of geographical origin were based on countries of origin.

The physical position of previously described *Pm* genes was either taken from the corresponding publication when available or estimated by blasting the flanking markers using BLASTN against Chinese Spring RefSeq v1.0 as a reference.

Accessions that possibly carry the causal genes of a resistance-associated region were determined by 1) having at least 50% of the resistance-associated SNPs within an associated region and 2) being resistant to the respective isolate (<=20% leaf coverage).

## Results

We assembled a geographically diverse panel of 755 bread wheat accessions (Fig 1a-c) based on the selective reduction of former collections used for allele mining of the powdery mildew resistance gene *Pm3* (Bhullar et al. 2009, 2010b). In the selection process, we focused on landraces and combined data on geographical origin with phenotypes of powdery mildew seedling stage resistance to six *Bgt* isolates. The resulting panel, hereafter LandracePLUS panel, contains 521 winter wheat and 234 spring wheat accessions, including 576 landraces, 162 older cultivars (acquisition date from 1946 to 2003, with the main part from the 60s and 70s), seven research or breeding lines and eleven unknown accessions (Table S1). We used the LandracePLUS panel, which covers a broad geographical distribution, with a focus on accessions originating from the Middle East (Fig 1a-b), to detect genetic loci associated with resistance to the powdery mildew pathogen (Fig 1d).

**Figure 1.**
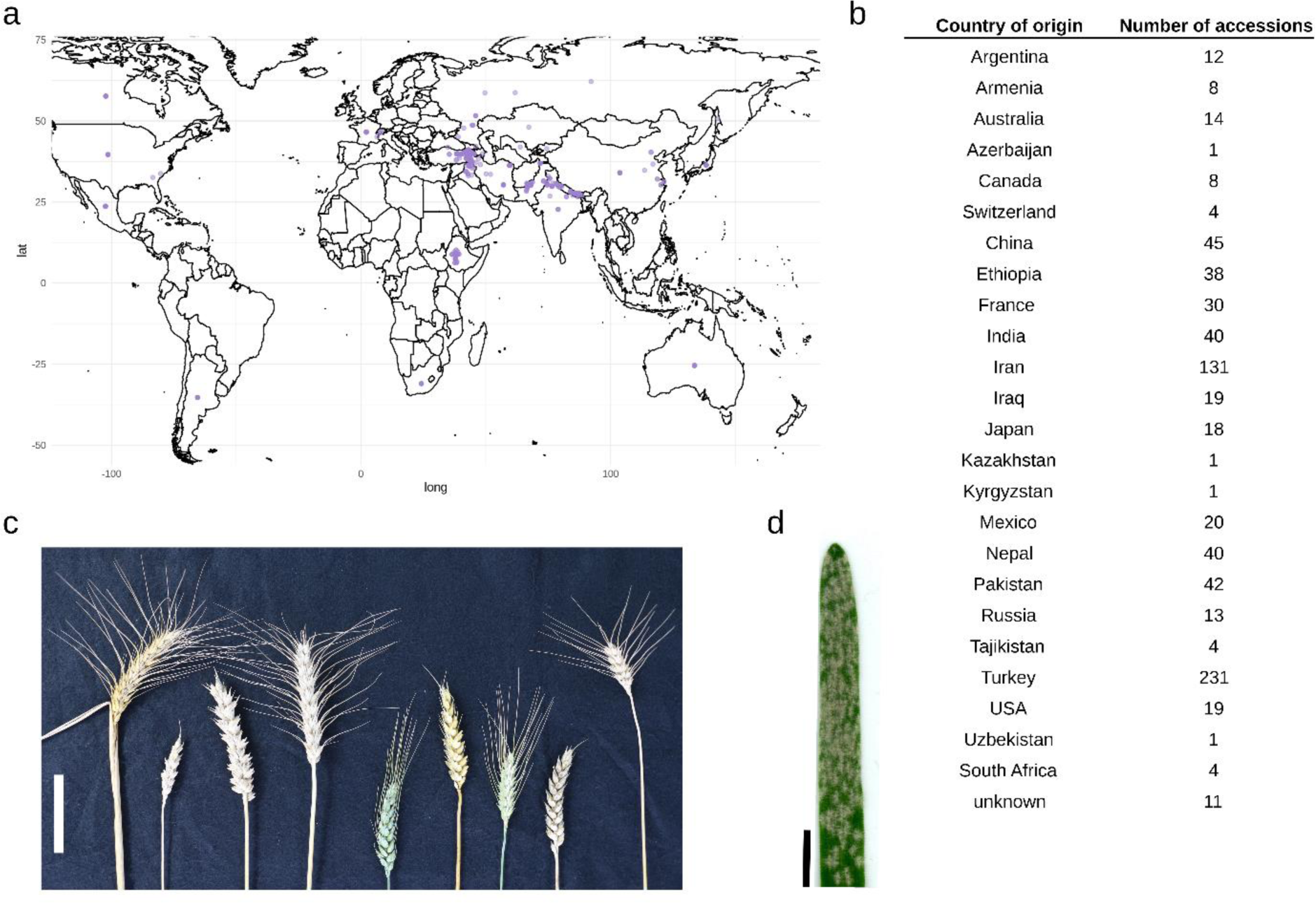
LandracePLUS panel diversity and powdery mildew symptoms. **a** World map with semitransparent purple dots representing the origin of each of the 755 wheat accessions of the LandracePLUS panel. **b** Number of accessions from the LandracePLUS panel per country of origin. **c** Selection of diverse wheat ears from the LandracePLUS panel. Scale bar, 5 cm. **d** Leaf of wheat cultivar “Kanzler” with powdery mildew symptoms. Scale bar, 1 cm

### Diversity analysis of the LandracePLUS panel reveals unexplored genetic diversity

A total of 29,965 high-quality, polymorphic SNPs derived from genotyping with the TaBW35K SNP array were used for diversity analysis. All these SNPs had known chromosomal positions based on the Chinese Spring reference genome assembly RefSeq v1.0 (IWGSC 2018), including 18,610 regular SNPs and 11,355 OTVs, i.e. markers that detect both presence-absence polymorphisms and nucleotide polymorphisms. The 29,965 markers were distributed across the wheat genome, similar to earlier findings (Liu et al. 2017; Alemu et al. 2021; Govta et al. 2022), with lower coverage of genome D compared to the A and B genomes: 11,684 (39.0%) markers on the A-genome, 13,589 (45.3%) on the B-genome and 4,692 (15.7%) on the D-genome. On average, 1,427 markers were assigned per chromosome, resulting in an average density of one marker each 483 Kbp. The least markers were assigned to chromosome 1D and most to chromosome 2B, with 497 and 2,465 markers, respectively.

For a better interpretation of the observed genetic diversity within the LandracePLUS panel, wheat accessions with high-quality genome sequences, namely the 10+ wheat reference genomes (IWGSC 2018; Walkowiak et al. 2020), Fielder (Sato et al. 2021) and Renan (Aury et al. 2022) were included in the analysis with 27,337 out of the 29,965 SNPs that mapped unambiguously to chromosomes of these genomes. Most accessions with reference genomes available clustered in Group 2 of the LandracePLUS panel, while Norin61 and Chinese Spring were on the edge of Group 4 (Fig 2a). A PCA (Fig 2a) and a dendrogram based on hierarchical clustering analysis (Fig 2b) revealed four genetic clusters correlated with geographical origin overall. Group 1 was dominated by accessions from Iran and Pakistan, while Group 2 was composed of accessions of mixed origin (Ethiopia, South Africa, USA, Canada, Mexico, Argentina, France, Switzerland, Kazakhstan, Kirgizstan, Tajikistan, Uzbekistan, Armenia and Russia). Group 3 predominantly contained accessions from Turkey, and Group 4 was dominated by South, Southwest, and East Asia (Iraq, Azerbaijan, India, Nepal, China and Japan).

**Figure 2.**
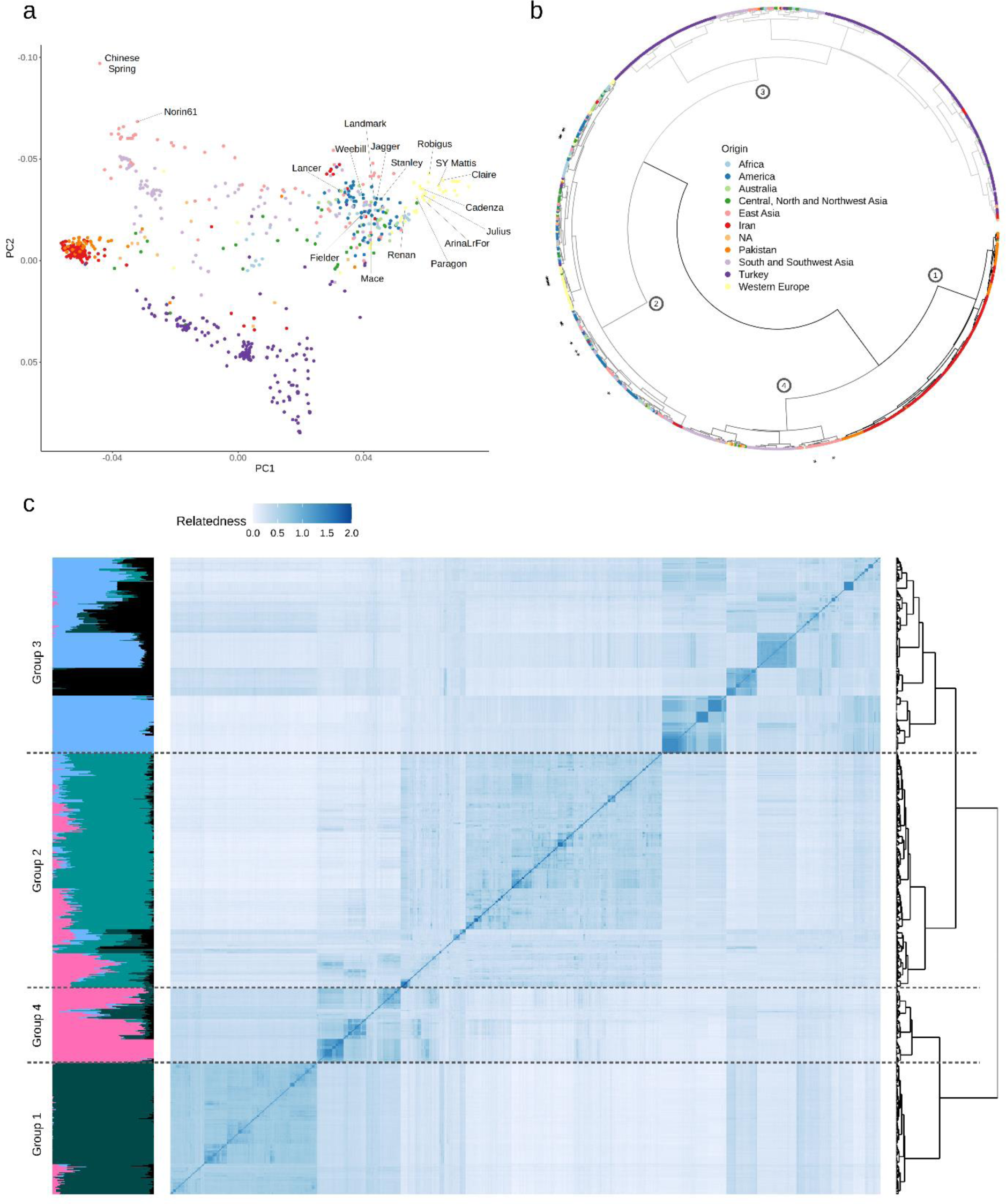
Genetic diversity and kinship analysis of the LandracePLUS panel. **a** PCA from 27,337 SNPs including high-quality sequenced wheat accessions labelled in the figure. PC1 = 8.4%, PC2 = 5.1%. Colors refer to the geographical origin of accessions and are indicated in b. **b** Dendrogram of a hierarchical clustering analysis from 27,337 SNPs including high-quality sequenced accessions indicated with stars. Colors represent the geographical origin of wheat accessions. Circled numbers on nodes refer to Groups 1 to 4 when dividing into four clusters. **C** Alignment of Admixture plot, kinship matrix and dendrogram for the 755 wheat accessions based on 29,965 SNPs. The Admixture plot shows K=5, where colors represent ancestral populations. In the kinship matrix, more saturated shades of blue indicate stronger relatedness. Dashed lines separate the four groups based on hierarchical clustering

When highlighting the different types of accessions in the PCA, landraces covered almost the entire genetic diversity range of the LandracePLUS panel. In contrast, the 162 cultivars clustered mainly with Group 2 and partially with Group 4 (Fig S2a). Therefore, landraces were genetically more diverse than cultivars and clustered apart from them. However, this was not always the case, as shown by the example of Ethiopian landraces, which clustered with cultivars in Group 2 (Fig S2b, Fig 2a), suggesting that Ethiopian landraces have substantially contributed to breeding programs. The close clustering of accessions from all over the world described above (Fig 2a-b) is likely driven by this separation between landraces and cultivars. As most of these geographically diverse accessions clustering together are cultivars, their genetic similarity is not derived from their origin but their breeding status as cultivars.

We assessed population structure performing hierarchical clustering on the 29,965 SNP set (excluding the high-quality genome sequences) and compared it to a kinship matrix and Admixture analysis (Fig 2c). The found clusters reflected the four groups described earlier, except for a shift of several accessions of diverse origin from Group 3 to Group 2. Estimated ancestral populations K=5 revealed a good fit for the LandracePLUS panel’s diversity into four groups (Fig S1). Yet, the kinship matrix revealed additional subdivisions within Groups 3 and 4.

To assess whether the LandracePLUS panel represents a similar genetic diversity compared to former studies on wheat germplasm, we compared our data with a collection of 632 landraces that were genotyped with the TaBW280K SNP array (Rimbert et al. 2018), which contains all markers that were used for the LandracePLUS panel (Balfourier et al. 2019). A PCA with 29,965 filtered SNPs (Fig S3) showed that the LandracePLUS panel covers the genetic diversity of this collection and further revealed unexplored genetic diversity absent in the study of Balfourier and colleagues (Balfourier et al. 2019). This additional diversity was mainly in Groups 1 and 3, comprising Turkish, Pakistani and Iranian landraces.

Taken together, the LandracePLUS panel is a diverse selection of wheat accessions with a pronounced diversity of landraces compared to cultivars and high-quality sequenced genomes. Furthermore, the panel covers earlier found genetic diversity and additionally expands it, mainly with Turkish, Pakistani and Iranian landraces.

### Differential lines reveal diverse virulence in a set of ten powdery mildew isolates

To maximize the chances of finding novel resistance genes, we first tested ten random isolates on a global collection of 27 differential lines, including NILs and donors of cloned or genetically described *Pm* genes (Fig 3). On average, isolates were avirulent on ten out of 27 differential lines. While isolates CHE_96224, CHE_97266 and CHE_98230 were avirulent on 16, 18 and 16 resistant differential lines, respectively, isolates CHE_19004 and ISR_94 were more virulent, with five and two resistant lines, respectively. Each isolate had a distinct virulence pattern, reflecting that the ten chosen mildew isolates represent broad diversity in virulence, confirmed by haplotype analysis of molecularly cloned *Avrs* (Table S3).

**Figure 3.**
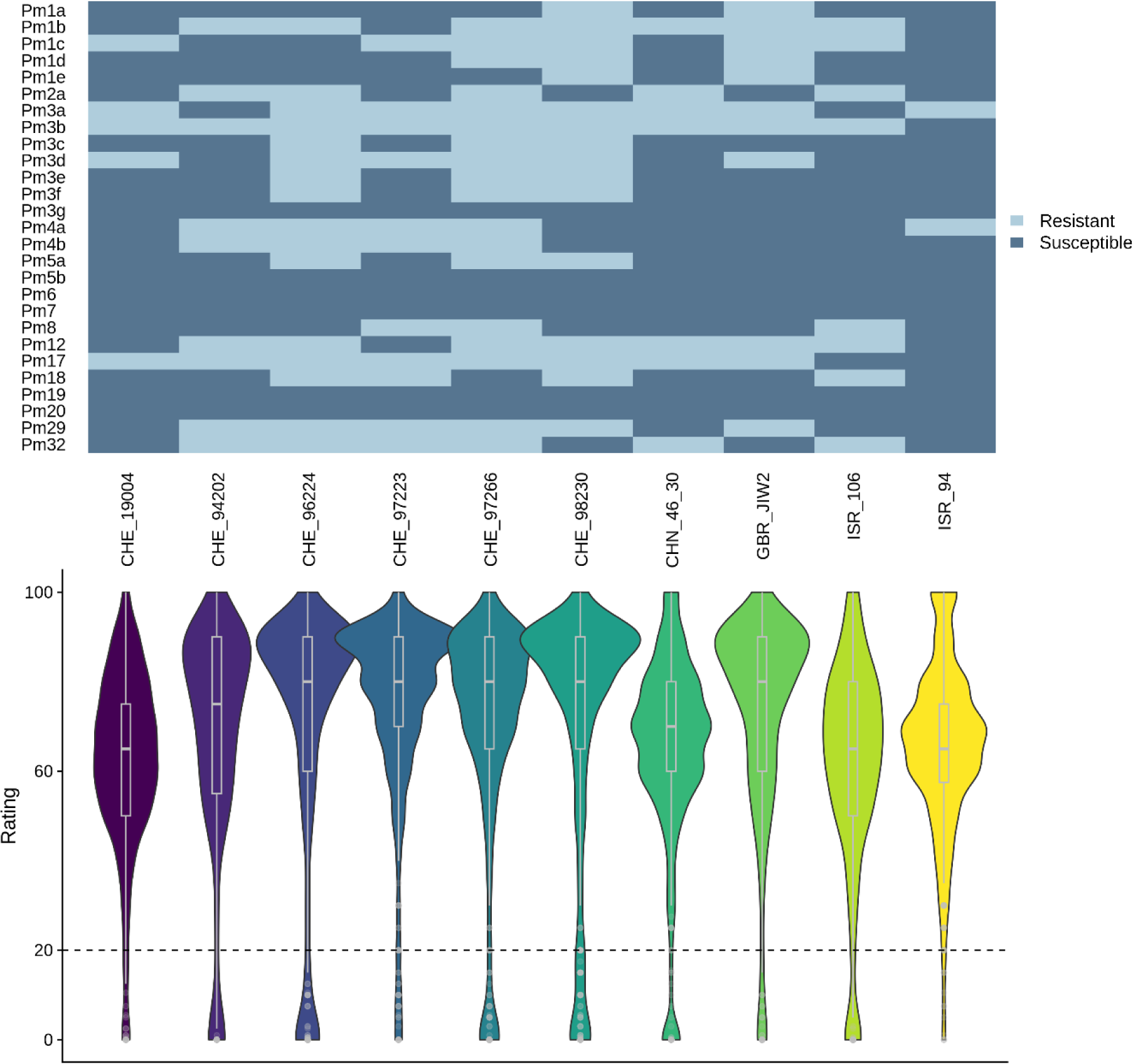
Virulence patterns of powdery mildew isolates and resistance distribution in the LandracePLUS panel. Virulence pattern of the ten powdery mildew isolates on differential lines with known *Pm* genes are shown as a heatmap on the top. On the bottom, the virulence pattern of the ten powdery mildew isolates on the LandracePLUS panel is depicted via violinplots and boxplots, highlighting median, 25 and 75 percentiles. The defined resistance threshold of 20 is indicated by a dashed line. Differential lines for the 27 *Pm* genes are listed in Table S2

Taken together, the resistance patterns of the differential lines reveal a phenotypically diverse set of ten powdery mildew isolates.

### The LandracePLUS panel shows varying resistance to ten wheat powdery mildew isolates

The response of the 755 wheat accessions to the ten powdery mildew isolates revealed an overall susceptibility, with an average of 56 resistant accessions (rating <=20) per isolate (Fig 3). Considering the response to individual isolates, only 35, 36, 40 and 17 wheat accessions were resistant to CHE_19004, CHE_97223, GBR_JIW2 and ISR_94, respectively. This reflected the broad virulence of CHE_19004 and ISR_94 already found in the differential lines, while CHE_97223 and GBR_JIW2 seemed to be more virulent on the LandracePLUS panel compared to the set of differential lines. On the other side, 91 and 89 wheat accessions were resistant to CHE_96224 and CHE_98230, respectively, which were the most avirulent isolates on the differential lines along with CHE_97266.

Phenotypic Pearson correlation between the isolates based on the set of differential lines corroborated the diversity of virulence (Fig S4a), with an average correlation coefficient of 0.31. The most similar isolates were CHE_96224 and CHE_97266, as well as CHE_98230 and GBR_JIW2, with a correlation of 0.69, while the two most diverging, though not significantly, were ISR_106 and ISR_94 with a negative correlation of 0.18. Pearson correlation between the isolates based on the LandracePLUS panel showed patterns resembling the differential line phenotypes (Fig S4b). While the overall relation between isolates was similar, several correlation coefficients differed notably, mostly involving ISR_106, GBR_JIW2 and CHE_98230. This deviation could be caused by a different content of *Pm* genes in the LandracePLUS panel compared to the differential set. With a coefficient of 0.36, however, the average correlation was similar. The diversity in resistance reactions observed within the LandracePLUS panel against these Bgt isolates suggests highly diverse effector content, highlighting the chances of finding novel resistance loci in the LandracePLUS panel.

No wheat accession showed resistance to all ten isolates, while nine wheat accessions were resistant to nine of the isolates. On average, excluding accessions susceptible to all tested isolates, wheat accessions were resistant to three isolates.

We predicted the presence of known *Pm* genes in accessions of the LandracePLUS panel that matched the pattern of the corresponding *Pm*-containing differential line. For example, 23 accessions of the LandracePLUS panel had the same resistance pattern as the *Pm2a* differential line and, hence, are good candidates for containing *Pm2a*. Indeed, 15 of these 23 accessions contained *Pm2* according to haplotype-specific markers (Manser et al. 2021) (Table S1). However, the potential presence of several *Pm* genes in the same accession can mask the resistance pattern of a specific *Pm* gene. This limits the use of differential lines to analyze overlapping resistance patterns and to predict specific genes. Nevertheless, the information on resistance patterns can be a useful tool to narrow down candidate accessions for the presence of a *Pm* gene of interest.

### The LandracePLUS panel shows isolate-dependent heritability of mildew response phenotypes and low correlation between phenotype and genetic relatedness

We calculated the phenotypic heritability using a linear mixed model approach. The effect of the wheat genotype on mildew resistance was isolate-dependent, between 0.38 for ISR_94 and 0.79 for CHE_96224, and with a batch effect of 0.48 and 0.12, respectively (Table S4). This suggests that the observed response to CHE_96224 is highly reproducible, while the batch seemed to have a strong influence on resistance reaction for ISR_94. Half of the isolates (CHE_94202, CHE_96224, CHE_97223, CHE_97266 and GBR_JIW2) had a heritability of 0.7 or higher, while ISR_94 was the only mildew isolate with a value below 0.5, possibly indicating a mixture of powdery mildew races. Accordingly, we removed isolate ISR_94 from all subsequent analyses. To account for this batch effect for the remaining isolates, and since the described nature of *R* genes can be considered a binary one – resistant or susceptible – we decided to transform the phenotypic scoring values of 0 to 100 to these two categories, with a threshold of 20 or higher for susceptibility. Thus, differences between batches are weighted less, providing more reliable phenotypes and GWAS results.

Using these categorized phenotypes, we further tested the correlation between the genetic relatedness and the phenotype using a Mantel test (Mantel and Valand 1970). There were only slight differences between the ten different isolates, and the correlations were close to zero, with Mantel r values ranging from -0.001 to 0.029, although not significant for most isolates (Table S5). This suggests that the applied approach of assembling the LandracePLUS panel minimized the effect of population structure on trait variation. The LandracePLUS panel should, therefore, provide improved power when conducting GWAS (Myles et al. 2009).

### Association studies for seedling resistance to wheat powdery mildew in the LandracePLUS panel reveal previously cloned *Pm* genes as well as possibly novel genes

We conducted GWAS with the phenotyping data obtained for each of the nine powdery mildew isolates on the LandracePLUS panel with a MAF of 1%. We first tested for a good fit of the univariate linear mixed model based on QQ plots, which was confirmed for all isolates except CHE_97223 (Fig S5). GWAS of the other eight isolates revealed five genomic regions associated with wheat mildew resistance on chromosomes 1A, 2B, 5D, 7A and 7D (Table 1, Fig S6). To account for LD, we defined a peak region by adding the average LD decay distance per subgenome from a recent study in wheat (Liu et al. 2023) to either side of the peak SNP. The most significant peak was located on chromosome 5D for the *AvrPm2*-containing *Bgt* isolates CHE_94202, CHE_96224, CHE_97266, CHN_46_30 and ISR_106. When focusing on one representative *AvrPm2*-containing isolate, ISR_106, the mildew resistance association spanned the region from 40,919,172 to 46,974,772 bp of the short arm of chromosome 5D of Chinese Spring (Fig 4a-c) and included the *Pm2* resistance gene locus (Sánchez-Martín et al. 2016). To test if the presence of *Pm2* was responsible for the significant association, we screened the LandracePLUS panel with a *Pm2* haplotype-specific marker (Manser et al. 2021) and found that out of 66 wheat accessions resistant to ISR_106, 31 contained *Pm2*. In the whole panel, 39 accessions contained *Pm2*, of which 34 were landraces (Table S1). Most of these 34 accessions were from Turkey, with three landraces from Russia, Pakistan and Tajikistan (Fig 4f). GWAS with a covariate for *Pm2* presence resulted in the loss of the significant peak (Fig 4d), corroborating that the peak was indeed caused by *Pm2*. Sequencing of the amplified *Pm2* locus in four randomly selected landraces that were positive for the haplotype marker uniformly revealed the presence of the known allele *Pm2a* (Sánchez-Martín et al. 2016).

**Table 1.**
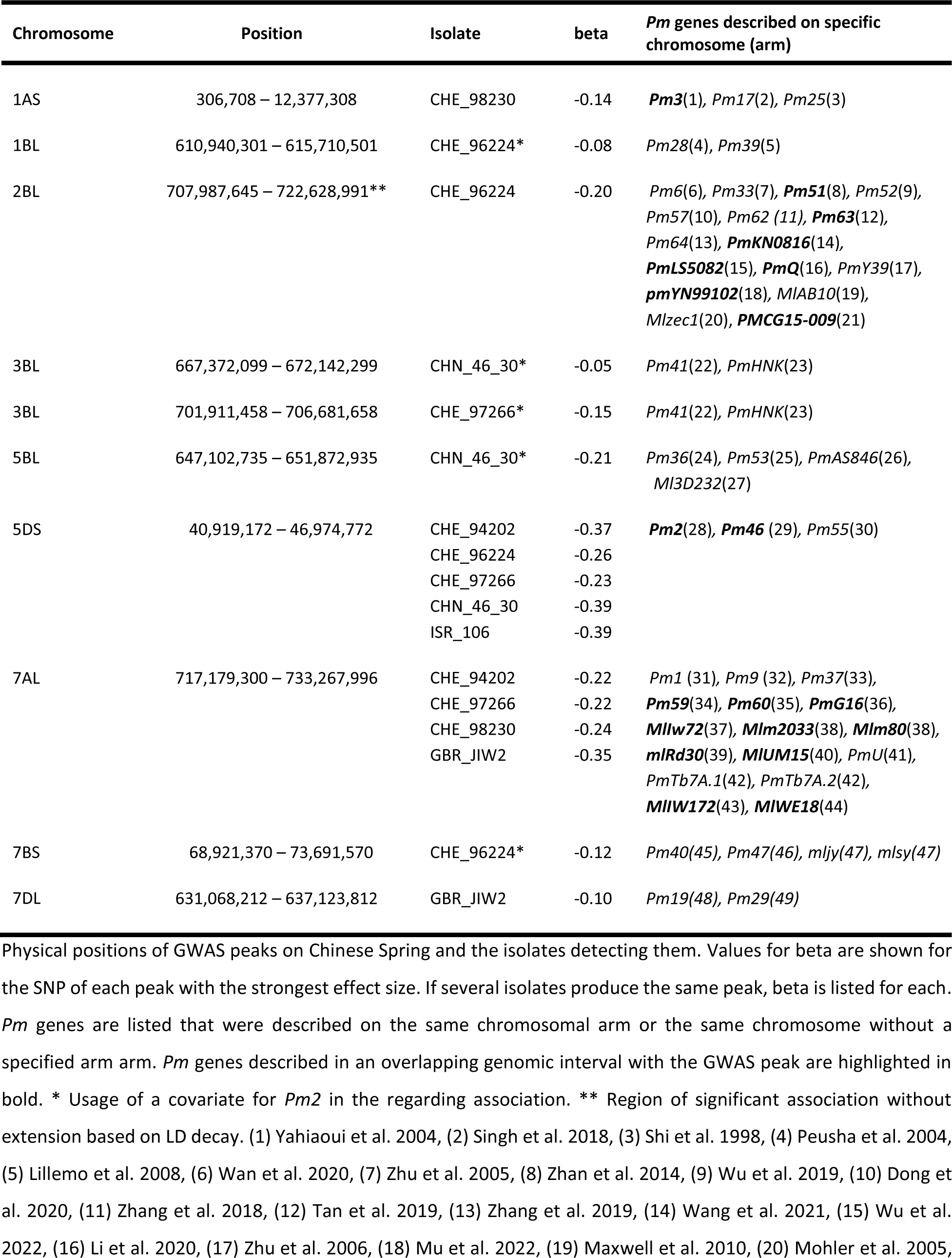
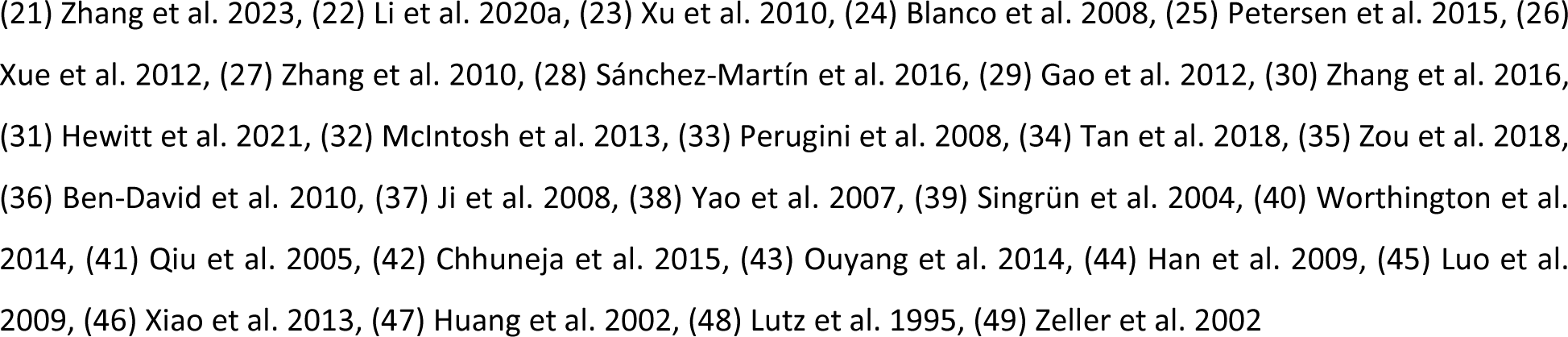
Powdery mildew associated regions detected in the LandracePLUS panel.

**Figure 4.**
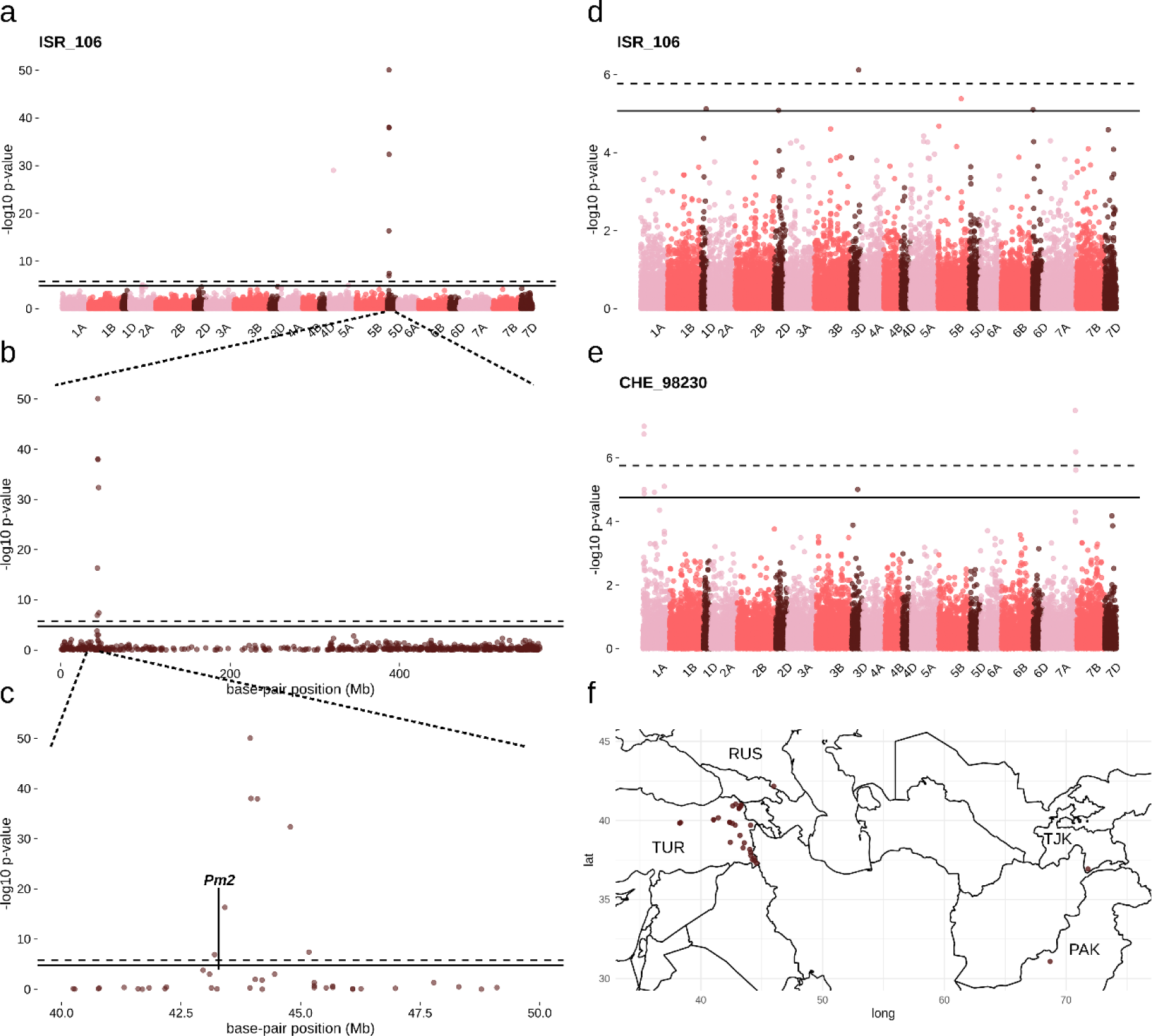
Analysis of the region associated with powdery mildew resistance on chromosome 5D. Manhattan plot for GWAS with all wheat accessions infected with isolate ISR_106 showing **a** all 21 chromosomes, **b** chromosome 5D and **c** the region of 40 to 50 Mb on chromosome 5D. The locus of the Chinese Spring version of *Pm2* with a partially deleted gene (Sánchez-Martín et al. 2016) is indicated with a line. **d** Manhattan plot for GWAS with ISR_106 when adding a covariate for *Pm2* presence in wheat accessions. **e** Manhattan plot for the *Pm2* virulent isolate CHE_98230. Solid lines represent the threshold for False discovery rate, and dashed lines for Bonferroni correction. **f** Map with semitransparent brown dots depicting the origin of landraces that contain *Pm2*. Countries of origin are abbreviated with the three-letter country code of ISO 3166

In addition to *Pm2*, we detected a peak on chromosome 1AS for CHE_98230, which spanned the genomic region from 306,708 to 12,377,308 bp (Table 1, Fig S6) and contains the locus of the cloned gene *Pm3*. Earlier studies have shown that out of 89 resistant accessions in the LandracePLUS panel, 21 contained functional *Pm3* alleles, with twelve and five accessions containing *Pm3c* and *Pm3b*, respectively (Table S1) (Bhullar et al. 2009, 2010a, 2010b). Therefore, we propose that the peak is caused by these functional *Pm3* alleles.

A third, very large region on chromosome 2BL significantly showed CHE_96224 resistance-associated SNPs from 707,987,645 to 722,628,991 bp (Table 1, Fig S6). Based on our criteria, this region was defined as three independent peaks. Upon closer inspection, the peaks are, however, only separated by small intervals showing no association. We therefore decided to consider them a single mildew-associated region. Indeed, it was shown that an introgression on chromosome 2BL is present in the wheat gene pool, most likely derived from the diploid wild relative *Triticum timopheevii* (Walkowiak et al. 2020; Keilwagen et al. 2022). This introgression is potentially present in the LandracePLUS panel. Many candidate *Pm* genes have already been described in this genomic interval, including *Pm51* (Zhan et al. 2014), *Pm63* (Tan et al. 2019), *PmKN0816* (Wang et al. 2021), *PmLS5082* (Wu et al. 2022), *PmQ* (Li et al. 2020b), *pmYN99102* (Mu et al. 2022) and *PmCG15-009* (Zhang et al. 2023).

We further detected a peak on chromosome 7AL for CHE_94202, CHE_97266, CHE_98230 and GBR_JIW2 that spanned from 717,179,300 to 733,267,996 bp (Table 1, Fig S6). Possible candidate genes in this genomic interval include *Pm59* (Tan et al. 2018), *MlIw72* (Ji et al. 2008), *Mlm2033* (Yao et al. 2007), *Mlm80* (Yao et al. 2007), *MlUM15* (Worthington et al. 2014), *MlIw172* (Ouyang et al. 2014), *PmG16* (Ben-David et al. 2010) and *mlRd30* (Singrün et al. 2004). Alleles of *Pm1* have also been mapped to chromosome 7AL (McIntosh et al. 2013). However, we could not confirm an overlap with our resistance-associated region because the *Pm1* locus is absent in Chinese Spring (IWGSC 2018).

Finally, we detected a peak on chromosome 7DL for GBR_JIW2 that spanned from 631,068,212 to 637,123,812 bp (Table 1, Fig S6, Fig 5a), where no *Pm* gene has been described previously.

**Figure 5.**
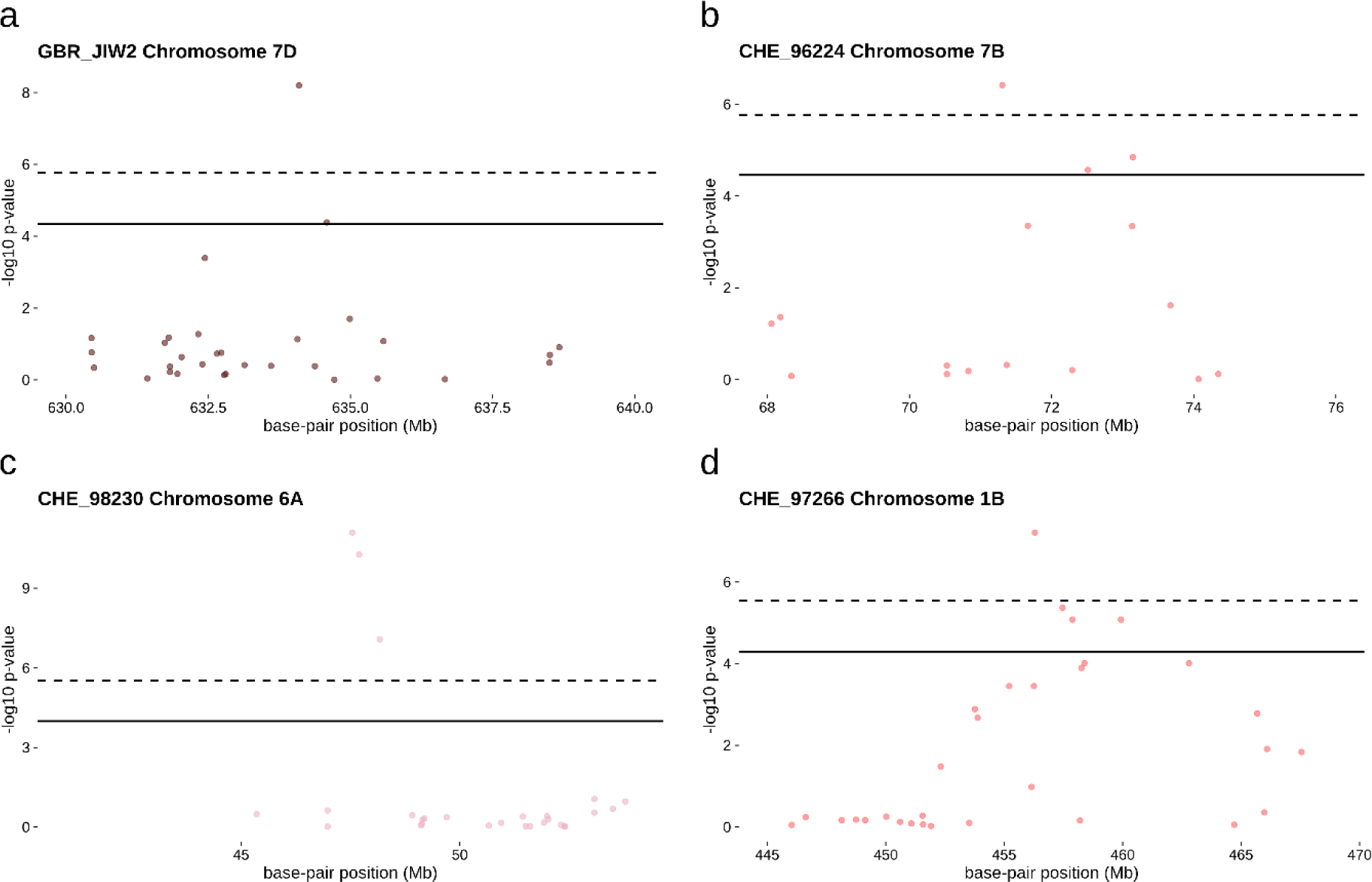
Manhattan plots of novel resistance associated regions. **a** All phenotyped LandracePLUS panel accessions. **b** All phenotyped LandracePLUS panel accessions using a *Pm2* covariate. **c** Subset of accessions from Pakistan and Iran. **d** Subset of accessions from Turkey with CHE_97266 representing the peak occurring for multiple isolates. Isolates and chromosomes are depicted in the top-left corner. Solid lines represent the threshold for False discovery rate and dashed lines for Bonferroni correction

### Association studies with a *Pm2* covariate reveal five additional and novel resistance-associated loci

We used the information on the presence of *Pm2* in LandracePLUS panel accessions to detect further resistance associations masked by the *Pm2* gene. Incorporating the covariate in GWAS for isolates with the *Pm2* peak, we detected five additional associations on chromosomes 1B, 3B, 5B and 7B (Table 1, Fig S7, Fig 5b). Two of these peaks appeared for CHE_96224, on chromosome 1BL, spanning from 610,940,301 to 615,710,501 bp and on chromosome 7BS from 68,921,370 to 73,691,570 bp. CHE_97266 showed an association on chromosome 3BL from 701,911,458 to 706,681,658 bp and the last two associations belonged to CHN_46_30, with a peak on chromosome 3BL from 667,372,099 to 672,142,299 bp and another on chromosome 5BL from 647,102,735 to 651,872,935 bp. These associations occur in regions where no *Pm* gene has been reported earlier, highlighting the utility of using a known *Pm* gene as a covariate to discover new resistance loci in GWAS.

### *Pm4* alleles are widely present in the LandracePLUS panel but are not revealed by association studies

Based on the resistance of differential lines carrying genes *Pm4a* or *Pm4b* to five and four of the nine isolates, respectively (Sánchez-Martín et al. 2021), we expected to detect the cloned gene *Pm4* in the LandracePLUS panel. Haplotype-specific markers detected the presence of the *Pm4* haplotype in 62 accessions. However, no peak was produced at the *Pm4* locus on chromosome 2AL for any isolate. To explain this missing association despite the broad presence of *Pm4*, we sequenced the locus in all 62 accessions, of which 51 and four accessions contained the non-functional alleles *Pm4f* and *Pm4g*, respectively (Sánchez-Martín et al. 2021; x), and four accessions with an undescribed allele, hereafter *Pm4_42460*, while the functional alleles *Pm4b* and *Pm4d* only occurred once and twice, respectively. The undescribed allele resembled *Pm4f*, except for one amino acid change in position 421 of splicing variant V1 (L421P). While allele *Pm4g* was present in accessions of diverse geographical origin, 42 out of 51 accessions carrying *Pm4f* and all four accessions containing *Pm4_42460* were Turkish landraces. Our findings suggest that the non-functional alleles *Pm4_42460* and *Pm4f* originated in Turkey.

Taken together, we find the presence of known *Pm* genes with GWAS of the LandracePLUS panel, giving insights into their geographic distribution and potential origin. In addition, we discovered six genomic regions where no *Pm* gene had been described earlier.

### Utilizing subsets of the LandracePLUS panel discovers novel loci that are associated with powdery mildew resistance of distinct geographical origin

Some resistance genes are likely present only in certain groups of the LandracePLUS panel, e.g., accessions with similar geographical origins like for the *Pm2* resistance gene. Accordingly, we first tested if subsets of the panel based on such groups were suitable to detect additional genomic regions associated with powdery mildew resistance. To assess the sensitivity of this approach, we used a re-sampling approach by creating 100 random subsets of 300, 200, 150 and 90 accessions from the LandracePLUS panel as input for GWAS with the *Pm2* avirulent isolate ISR_106. Since the MAF threshold of 1% used above would not filter out SNPs that occur only twice for a subset of 200 to 299 accessions, we adjusted the MAF threshold to 5% for all subsets below a sample size of 300. We found the *Pm2*-associated peak in 100%, 87%, 78% and 52% of cases for the four subset sizes, respectively. Therefore, we considered subsets of a minimum of 150 to 200 accessions suitable for detecting loci associated with a major resistance gene in the LandracePLUS panel.

In our sub-setting approach, we created sets of accessions based on their geographical origins. A subset of landraces from Pakistan and Iran showed a good fit to the GWAS model only for the mildew isolate CHE_98230 (Fig S8), revealing five additional peaks compared to the full LandracePLUS panel, located on chromosomes 2B, 5A, 5B, 5D and 6A (Table 2, Fig S9, Fig 5c). While the peak on chromosome 2BL was located in a genomic region that has been described to contain the resistance gene *Pm51* (Zhan et al. 2014) and the peak on chromosome 5BL has been implicated earlier with the resistance genes *Pm53* (Petersen et al. 2015) and *Ml3D232* (Zhang et al. 2010), the associated regions on chromosomes 5AL from 679,959,567 to 692,030,167 bp and on 5DS from 26,487,899 to 32,543,499 bp have not been associated with powdery mildew resistance before to our knowledge (Table 2). The last resistance-associated region, on chromosome 6AS, spanned the region from 41,508,045 to 53,578,645 (Fig 5c). On chromosome arm 6AS, only *Pm21* and *Pm56* have been described. *Pm21* originated and was cloned from the diploid grass *Dasypyrum villosum* (He et al. 2018; Xing et al. 2018). The gene was introduced in the hexaploid gene pool in China through a translocation line T6AL.6VS in the late 1980s (Chen et al. 1995). However, the translocated arm from *D. villosum* does not recombine with wheat homeologs (He et al. 2017). Hence, we expect to see a broad association covering the short arm of chromosome 6A in the case of *Pm21* detection. Further, *Pm21* was described to confer broad-spectrum resistance (He et al. 2017), whereas we observed the association for only one out of ten isolates. Therefore, we assume that the causal gene in our resistance-associated region differs from *Pm21*. Similarly, *Pm56* was only recently introduced as a translocation line 6AL.6RS from rye (Hao et al. 2018) several years after we obtained and utilized the seeds of the initial collection. We conclude that this is a resistance-associated region on chromosome 6AS not previously described.

**Table 2.** Powdery mildew associated regions detected in subsets of the LandracePLUS panel.

| Chr | Position | Subset | Isolate | beta | *Pm* genes described on specific chromosome (arm) |
| --- | --- | --- | --- | --- | --- |
| 1AL | 539,831,968 – 551,902,568 | Turkey (221) | CHE_98230 | -0.14 | *Pm25*(1) |
| 1BL | 453,908,676 – 462,314,398 | Turkey (223, 222, 222, 217, 212) | CHE_94202<br>CHE_96224<br>CHE_97266<br>CHN_46_30<br>ISR_106 | -0.22<br>-0.24<br>-0.26<br>-0.28<br>-0.26 | *Pm28*(2), *Pm39*(3) |
| 1BL | 664,434,720 – 669,204,920 | Turkey (222) | CHE_96224 | -0.22 | *Pm28*(2), *Pm39*(3) |
| 1DL | 333,346,020 – 339,401,620 | Turkey (217, 212) | CHN_46_30<br>ISR_106 | -0.48<br>-0.48 | *Pm10*(4) |
| 2AL | 534,734,122 – 546,804,722 | Turkey (221) | CHE_98230 | 0.38 | *Pm4*(5), *Pm50*(6), *Pm65*(7), *pmX*(8), *PmHo*(9), *MI92145E8-9*(10), *PmHnk54*(11), *PmLK906*(12), *PmYm66*(13), *PmSN15218*(14) |
| 2AL | 762,574,656 – 774,645,256 | Turkey (221) | CHE_98230 | 0.84 | *Pm4*(5), *Pm50*(6), ***Pm65***(7), ***pmX***(8), *PmHo*(9), *MI92145E8-9*(10), *PmHnk54*(11), *PmLK906*(12), *PmYm66*(13), ***PmSN15218***(14) |
| 2BL | 737,551,805 – 742,322,005 | Pakistan/Iran (167) | CHE_98230 | -0.33 | *Pm6*(15), *Pm33*(16), ***Pm51***(17), *Pm52*(18), *Pm57*(19), *Pm62* (20), *Pm63*(21), *Pm64*(22), *PmKN0816*(23), *PmLS5082*(24), *PmQ*(25), *PmY39*(26), *pmYN99102*(27), *MIAB10*(28), *MIzec1*(29), *PMCG15-009*(30) |
| 3AL | 643,922,092 – 655,992,692 | Turkey (220) | GBR_JIW2 | -0.16 | - |
| 4BL | 610,988,411 – 615,758,611 | Turkey (224) | CHE_97223 | -0.12 | *Pm7*(31), *Mld*(32), *PmTx45*(33) |
| 5AL | 472,287,696 – 484,857,509 | Turkey (217) | CHN_46_30 | 0.16 | *pm2026*(34) |
| 5AL | 679,959,567 – 692,030,167 | Pakistan/Iran (167) | CHE_98230 | 0.28 | *pm2026*(34) |
| 5BL | 375,685,467 – 380,455,667 | Turkey (223) | CHE_94202 | -0.20 | *Pm36*(35), *Pm53*(36), *PmAS846*(37), *MI3D232*(38) |
| 5BL | 517,686,966 – 522,457,166 | Pakistan/Iran (167) | CHE_98230 | 0.19 | *Pm36*(35), ***Pm53***(36), *PmAS846*(37), ***MI3D232***(38) |
| 5BL | 656,831,925 – 661,602,125 | Turkey (217) | CHN_46_30 | 0.19 | *Pm36*(35), *Pm53*(36), *PmAS846*(37), *MI3D232*(38) |
| 5DS | 26,487,899 – 32,543,499 | Pakistan/Iran (167) | CHE_98230 | 0.27 | *Pm2*(39), *Pm46* (40), *Pm55*(41) |
| 6AS | 0 – 7,996,136 | Turkey (224) | CHE_97223 | -0.12 | *Pm21*(42, 43), *Pm56*(44) |
| 6AS | 41,508,045 – 53,578,645 | Pakistan/Iran (167) | CHE_98230 | -0.25 | *Pm21*(42, 43), *Pm56*(44) |
| 7AL | 611,542,221 – 623,612,821 | Turkey (224) | CHE_97223 | -0.09 | *Pm1*(45), *Pm9* (46), *Pm37*(47), *Pm59*(48), *Pm60*(49), *PmG16*(50), *Mllw72*(51), *Mlm2033*(52), *Mlm80*(52), *mIRd30*(53), *MIUM15*(54), *PmU*(55), *PmTb7A.1*(56), *PmTb7A.2*(56), *Mllw172*(57), *MIWE18*(58) |
| 7BL | 701,729,427 – 706,653,524 | Turkey (221) | CHE_98230 | -0.16 | ***Pm5***(59), ***pmDHT***(60), ***PmSGD***(61),<br>***PmTm4***(62), ***pmHYM***(63), ***pmYBL***(64),<br>***mlxbd***(65), *mljy*(66), *mlsy*(66) |
| 7DL | 497,931,007 – 503,986,607 | Turkey (223,<br>222, 217, 212) | CHE_94202<br>CHE_96224<br>CHN_46_30<br>ISR_106 | -0.16<br>-0.15<br>-0.16<br>-0.19 | *Pm19*(67), *Pm29*(68) |

Finally, we used the same geographical approach to investigate only Turkish landraces, revealing a good fit to the model for all isolates except CHE_19004 (Fig S10). This subsetting resulted in the discovery of 15 additional peaks on chromosomes 1A, 1B, 1D, 2A, 3A, 4B, 5A, 5B, 6A, 7A, 7B and 7D (Table 2, Fig S11). In earlier studies, two of the 15 regions have been described with powdery mildew seedling resistance. The associated region on chromosome 2AL from 762,574,656 to 774,645,256 bp covers three known *Pm* genes, while the peak on chromosome 7BL from 701,729,427 to 706,653,524 bp overlaps with eight previously described *Pm* genes (Table 2). The association on chromosome 1BL from 664,434,720 to 669,204,920 bp includes the *Pm39* locus. However, *Pm39*, also known as *Lr46* (Lillemo et al. 2008), is an adult plant resistance gene and not active at seedling stage, and we conclude that the detected association is caused by an unknown, novel gene. The remaining twelve loci do not overlap with previously described *Pm* genes and therefore depict good candidates for novel powdery mildew resistance loci (Table 2).

Taken together, subsets based on geographical origin revealed 16 genomic regions where no *Pm* genes have been described previously, suggesting that these genes arose in the respective countries Pakistan, India and Turkey.

### Candidate genes of five novel resistance loci include putative NLRs, serine/threonine kinases, a C2H2-type zinc finger and F-box-like proteins with leucine-rich repeat (LRR) domains

We investigated five of the 22 novel resistance loci more closely to get an insight into possible candidate genes (Figure 5, Table S6). We chose the most significant peak with at least two significantly associated SNPs from GWAS of 1) the full LandracePLUS panel, 2) the panel using the *Pm2* covariate and 3) the Pakistan/Iran subset, on chromosomes 7DL, 7BS and 6AS, respectively. From the subset of Turkish landraces, we chose the highly significant single SNP association on chromosome 1DL and the peak on chromosome 1BL that occurred for five isolates, suggesting a more broad-spectrum resistance. For the associated region on chromosome 7DL, candidates annotated on Chinese Spring were ten putative NLRs, while the associated region on chromosome 7BS contained an annotation for a C2H2-type zinc finger and one F-box-like protein with an LRR domain. All associated regions except for 1BL had annotations for one serine/threonine kinase, the only candidate for the peak on chromosome 1DL. Both peaks on chromosome 6AS and 1BL each contained annotations for one putative NLR and the peak on chromosome 6AS included four additional F-box-like proteins with LRR domains (Figure 5, Table 6). Further validation studies are necessary to confirm the resistant nature of these candidate genes.

### Novel resistance loci for breeding programs

We analyzed the gene pool of the twenty most modern cultivars of the LandracePLUS panel (registered between 1990 and 2003) to evaluate whether the 22 potentially novel powdery mildew genes are present in elite material or potentially novel in this gene pool (Supplementary_file3). For genomic regions on chromosomes 5BL (derived from the LandracePLUS panel using a *Pm2* covariate) and 6AS (from the Pakistan/Iran subset), the most modern cultivars contained less than 50% of the resistance-associated SNPs, suggesting that the underlying resistance genes are likely not present in these cultivars. For six of the 22 associations, including the peaks on chromosomes 7DL and 1BL (described above in detail), some of the cultivars contained at least 50% of the resistance-associated SNPs, but none showed resistance towards the respective mildew isolate. We conclude that the underlying resistance genes are not present in this germplasm, at least not as active, resistance-conferring alleles. These findings suggest that the resistance loci have not been transferred to the modern gene pool from landraces. However, investigation of more recent cultivars would be needed to confirm these results.

Resistance-associated SNPs of the remaining 14 associated regions, including the peaks on chromosomes 7BS and 1DL discussed extensively, were present in the most modern germplasm. However, few accessions that harbored the alleles were resistant. Thus, the underlying resistance-conferring alleles seem to be partially present, meaning that breeders could integrate the corresponding cultivars we highlighted (Supplementary_file3) directly in their programs, avoiding possible yield penalties due to linkage drag. However, the combination of resistance and presence of resistance-associated SNPs occurred only for the three cultivars TRI 17181, TRI 17284 and TRI 16947. While these cultivars are attractive resistance-breeding candidates, it is difficult to dissect which detected regions are causing the observed resistance and, therefore, actually contain causative alleles.

## Discussion

### The LandracePLUS panel harbors untapped genetic diversity originating mainly from Turkish, Pakistani and Iranian landraces

We assembled a diverse panel of 755 bread wheat accessions with a focus on landraces. A FIGS approach laid the foundation for a trait-customized collection of accessions with potentially high selection pressure for powdery mildew resistance. A second step of selective reduction based on geographical origin resulted in the LandracePLUS panel. Mantel tests revealed a small impact of the population structure on powdery mildew resistance variation, reflecting the successful outcome of our targeted panel assembly. The resulting improved power of GWAS (Myles et al. 2009) facilitated the detection of novel *Pm* genes in the LandracePLUS panel.

The LandracePLUS panel revealed untapped genetic diversity in Turkish, Iranian and Pakistani landraces. It showed four main genetic clusters that correlate with the geographical origin, similar to a study which showed that wheat accessions from the Caucasus region, as well as Central, South and East Asian are more diverse compared to other regions in the world (Balfourier et al. 2019). Landraces covered almost the full diversity of the LandracePLUS panel, whereas cultivars were limited to one cluster. This cluster also included the bread wheat accessions with high-quality sequenced genomes, except for the two landraces Chinese Spring and Norin61 (IWGSC et al. 2018; Walkowiak et al. 2020; Sato et al. 2021; Aury et al. 2022). This observation highlights the need to include landraces in breeding programs to enlarge the genetic base of elite wheat varieties (Lopes et al. 2015; Marone et al. 2021). Future diversity studies of wheat should focus on landraces, in particular on those from regions with untapped genetic diversity, such as Turkey, Pakistan and Iran.

### Pathogen virulence characterization guides *Pm* gene discovery

To discover novel *R* genes against powdery mildew in the LandracePLUS panel, we used a set of ten *Bgt* isolates that showed highly diverse virulence patterns when tested on differential lines with single, known *Pm* genes. As we observed avirulence for many differential lines, using these isolates should reveal the presence of most *Pm* genes for which differential lines are available.

The tested wheat accessions were rarely resistant to more than three isolates, in line with characteristic *R* gene-based race-specific resistance (Flor 1971). This implies that the observed single resistance genes would be of limited agricultural use, depending on the isolates present in the corresponding wheat-growing area. To broaden the resistance spectrum and prevent fast evolution of pathogen virulence, such genes should be deployed in a suitable manner, e.g., by gene stacking, gene pyramiding or transgenic overexpression (Mundt 2018; Koller et al. 2019). On the other hand, the accessions of the LandracePLUS panel susceptible to all tested powdery mildew isolates can be assumed to lack any major resistance gene active at the seedling stage. Thus, observed resistance in adult plants in the field would likely be durable, making such accessions attractive donors of adult plant resistance.

Currently, for the 27 genes represented in the differential set of *Pm* lines used in this study, only eight of the corresponding avirulence genes are molecularly known (Bourras et al. 2015, 2019; Praz et al. 2017; Hewitt et al. 2021; Müller et al. 2022; Kloppe et al. 2023; Kunz et al. 2023). Avirulence gene sequence comparison in our ten powdery mildew isolates revealed the presence of recognized haplotypes of seven of the cloned Avrs. This knowledge guided us to determine whether a resistance-associated region in the LandracePLUS panel was derived from a known *Pm* gene, as we showed for the cloned gene *Pm2a*. Further, the information on resistance to *Bgt* isolates of distinct geographical origin can guide an informed deployment of accessions in the respective agricultural areas (Vleeshouwers and Oliver 2014; Müller et al. 2022). For example, our findings suggest further work on landraces resistant to CHN_46_30 for potential deployment in China because they must contain effective *R* genes against this isolate of Chinese origin.

### GWAS of the LandracePLUS panel detects known *Pm* genes and reveals six undescribed powdery mildew resistance-associated regions on chromosomes 1BL, 3BL, 5BL, 7BS and 7DL

GWAS for eight powdery mildew isolates on the LandracePLUS panel without subsetting revealed ten resistance-associated regions. Four of these were in genomic regions with previously described resistance genes. Of these, the peaks on chromosomes 2BL and 7AL overlapped with regions of genetically described, but not molecularly known *Pm* genes, while the peaks on chromosomes 1AS and 5DS were likely caused by the cloned genes *Pm3* and *Pm2*, respectively. While *Pm3* was known to be present in several LandracePLUS panel accessions (Bhullar et al. 2009, 2010a, 2010b), we did not expect to find *Pm2* widely in the gene pool of landraces. *Pm2* was introduced into the breeding gene pool via the Russian cultivar Ulka (Pugsley and Carter 1953). It originated from the diploid wheat wild relative *Ae. tauschii* and has eight known haplotypes, of which only *Pm2a* was detected in hexaploid wheat (Manser et al. 2021). While *Pm2a* has been identified previously in six wheat landraces (Chen et al. 2019; Manser et al. 2021), we found its presence in 34, mostly Turkish landraces, suggesting Turkey as the geographical origin of the *Pm2a* resistance gene. While GWAS did not detect the cloned gene *Pm4* in the LandracePLUS panel, we discovered a high frequency of two non-functional *Pm4* alleles in Turkish landraces. Our findings suggest that these alleles originated in Turkey, fitting the origin of *Pm4* from tetraploid wheat, which also arose in Turkey (Özkan et al. 2002; Sánchez-Martín et al. 2021). Despite the narrow representation of *Pm4b* and *Pm4d* in landraces, a wide presence of these functional alleles compared to *Pm4f* and *Pm4g* has been described in elite germplasm (Sánchez-Martín et al. 2021). This suggests that the breeding process for *Pm4* mildew resistance was very effective.

We detected one region on chromosome 7DL not previously described as associated with mildew resistance. This region contained annotations for ten putative NLRs and one serine/threonine kinase. While NLRs are to date still the most common candidates for *Pm* genes, serine/threonine kinases have been described to play a role in defense response, including *Pto*, which confers resistance to bacterial speck disease in tomato (Martin et al. 1993; Loh and Martin 1995).

Using a *Pm2* covariate, we detected five additional undescribed resistance associations in the LandracePLUS panel. One of them was located on chromosome 7BS, containing candidate genes putatively encoding a C2H2-type Zinc finger, a serine/threonine kinase and an F-box-like protein with LRR domains. The latter have been described to facilitate hypersensitive cell death response in tobacco and tomato (Van Den Burg et al. 2008) and shown to be involved in defense response to stripe rust in wheat (Yin et al. 2018). On the other hand, Zinc fingers of the C2H2-type were linked to plant defense response (Kim et al. 2004; Tian et al. 2010; Yin et al. 2020; Sharma et al. 2021), where some cases have shown that the transcriptional repression activity of the zinc finger was the mechanism behind this association (Weigel et al. 2005; Uehara et al. 2005).

### Targeted GWAS subsets reveal 16 potentially novel resistance-associated loci

Utilizing targeted subsets, we discovered 16 most likely novel peaks on ten chromosomes (Table 2). GWAS are expected to be more powerful when conducted on large datasets where individuals are drawn randomly from the population (Uffelmann et al. 2021). However, important SNPs at a small regional scale might yet be diluted in species-wide panels and not detected by GWAS, which typically lacks the power to detect associations with rare alleles (Marees et al. 2018). Pending sufficient phenotypic and genetic variation, GWAS in local panels have proven very effective in *Arabidopsis thaliana* (Gloss et al. 2022). For this reason, resistance genes which may have been selected at small regional scales might be more efficiently detected in subsets of accessions from the same geographical origin.

Therefore, we focused on accessions originating from distinct countries that harbored novel genetic diversity, namely Pakistan, Iran and Turkey. This resulted in the detection of 16 additional loci where no *Pm* genes have been described. With a subset of accessions exclusively from Pakistan or Iran, we discovered a region on chromosome 6AS associated with mildew resistance. The region contains genes putatively encoding an NLR, a serine/threonine kinase and F-box-like proteins with LRR domains.

While we present novel regions and gene candidates, the molecular nature of the observed resistance must be confirmed in future studies, especially when considering that the reference genome Chinese Spring might lack (susceptible) alleles of the causal resistance genes. One approach would be the application of recent sequencing technologies, such as circular consensus sequencing (CCS) (Wenger et al. 2019). Such novel approaches have increased the feasibility of sequencing single donor accessions to assist in the cloning a gene of interest. For example, assembling the Kariega genome with CCS has demonstrated the usefulness of this approach in wheat and has led to the cloning of *Yr27* (Athiyannan et al. 2022a). Another option to identify a gene of interest in a specific genotype depends on the availability of a pangenome, which ideally would capture the entire gene repertoire of a species (Tettelin et al. 2005). High-quality sequencing efforts have recently resulted in 19 wheat genomes, of which 14 have reference genome quality (IWGSC et al. 2018; Walkowiak et al. 2020; Sato et al. 2021; Aury et al. 2022; Athiyannan et al. 2022a; Kale et al. 2022). Genome analysis has revealed structural rearrangements, introgressions and differences in gene content (Walkowiak et al. 2020). A successful example of using this resource is the cloning of *Lr14a*, which was based on the reference genome ArinaLrFor (Kolodziej et al. 2021). However, despite the recent advances in pangenome projects, the close clustering of the high-quality sequenced genomes compared to the LandracePLUS panel suggests that the currently available pangenome includes only a fraction of the diversity present in landraces, including resistance loci. Thus, it is essential to include more diverse wheat accessions, specifically landraces, in future work to increase the extent of the pangenome, particularly for NLR loci, which are rarely present across wheat genotypes. We propose to assemble high-quality genomes of several landraces from Turkey, Pakistan and Iran to achieve such a goal. Furthermore, contrasting phenotypes for traits of interest should be included when setting up pangenome consortia. This diversified selection could provide a resource that guides various trait-genotype associations. Finally, we suggest choosing a donor accession with confirmed resistance for each peak and sequencing the resistance-associated genome using CCS. This will reveal which gene candidates are present in the specific accessions and guide their molecular cloning and validation, e.g., via virus-induced gene silencing (Cakir et al. 2010).

The subset-based association analysis done in this work of the genetically diverse LandracePLUS panel challenged with ten *Bgt* isolates unraveled 22 potentially novel powdery mildew resistance genes. Therefore, this study can be used as an example for future work on similar collections in search of other traits of interest. Once a diversity panel is assembled, instead of focusing on single pathogen races, phenotyping with diverse isolates from geographically defined agricultural regions followed by subset-based analyses for the origin of accessions, would reveal more resistance loci compared to studies done with single pathogen isolates on entire collections.

## Acknowledgments

We would like to thank Nikolaos Minadakis, Benjamin Jaegle, Zoe Bernasconi, Alexandros Sotiropoulos, Thomas Wicker, Bruno Studer and Anne Roulin for their input and guidance in acquiring necessary techniques for this study. We would further like to thank Sarah Furrler, Garp Linder and Pietro Sassi for their contributions to this project in the lab. This project was financially supported by the University of Zurich, Swiss National Science Foundation grants 310030_204165 and 310030B_182833 to B.K. JSM is recipient of the grants “Ramon y Cajal” Fellowship RYC2021-032699-I, funded by MCIN/AEI/10.13039/501100011033 and by the “European Union NextGenerationEU/PRTR” and PID2022-142651OA-I00, funded by MCIN/AEI/10.13039/501100011033 and “European Union NextGenerationEU/PRTR”.

## Statements & Declarations

### Funding

This project was financially supported by the University of Zurich, Swiss National Science Foundation grants 310030_204165 and 310030B_182833 to B.K. JSM is recipient of the grants “Ramon y Cajal” Fellowship RYC2021-032699-I, funded by MCIN/AEI/10.13039/501100011033 and by the “European Union NextGenerationEU/PRTR”, and PID2022-142651OA-I00, funded by MCIN/AEI/10.13039/501100011033 and by the “European Union NextGenerationEU/PRTR”.

### Competing Interests

Beat Keller is a member of the editorial board of TAG. The authors declare no other competing interests.

### Author contribution statement

J.S.M. and B.K. conceived the project. E.J. and R.L. performed seed propagation. R.L., V.W. and J.S.M. performed infection tests. R.L. performed DNA extraction, PCR screens and allele detection. E.P. performed SNP array genotyping. R.L. and M.H. performed bioinformatics analysis. R.L., J.S.M. and B.K. analyzed the data and wrote the manuscript. All authors revised the manuscript.

### Data Availability

The mildew isolate CHE_19004 whole-genome sequence data is available at NCBI’s Short Read Archive (SRA) under the accession code BioProject PRJNA945619. The nine isolate sequences from earlier work (Sotiropoulos et al. 2022) are available at NCBI under PRJNA625429 and SRP062198. The genotyping data of landraces from the study of Balfourier and colleagues (Balfourier et al. 2019) can be accessed at https://urgi.versailles.inra.fr/download/wheat/genotyping/Balfourier_et_al_Wheat_Phylogeography_D ataS2.zip. All other data needed to evaluate the conclusions in this study are included in the article and its supplementary information files. All *Blumeria graminis* f.sp. *tritici* isolates used in this study are kept alive in the Department of Plant and Microbial Biology of the University of Zurich and are available upon request.

## Supporting information captions

**Supplementary file1**

**Supplementary file2**

**Table S1. Information on accessions of the LandracePlus panel.** Alleles with asterisks are non-functional. Groups are based on the hierarchical clustering of the LandracePLUS panel, including high-quality sequenced genomes.

**Table S2. Differential lines used for determining powdery mildew isolate virulence spectra and the respective Pm genes they contain.**

NILs had been backcrossed multiple times with susceptible accessions “Federation” or “Chancellor”, depicted by /x*Accession, where x is the number of backcrosses to the designated accession (McIntosh et al. 2013).

**Table S3. Molecularly known Avr and Svr haplotypes of the ten powdery mildew isolates used for phenotyping.** H0 = published avirulent haplotype, H1 = published virulent haplotype, H2 = unpublished haplotype. Based on the phenotype of the differential lines containing cognate Pm genes, we propose that H2 of AvrPm1a and AvrPm3a/f are virulent, while H2 of AvrPm3d is proposed to be avirulent. H2 of AvrPm17 is likely virulent due to the resemblance of H1, with just one additional amino acid change. The resistant phenotype of the differential line containing Pm17 is suggested to be due to an additional Pm gene in Amigo. Amino acid (AA) changes are given based on H0 and depicted with original AA, position, new AA.

**Table S4. Heritability of phenotypes for all ten powdery mildew isolates based on a linear mixed model.**

Table S5. Correlation of genetic relatedness and phenotypes for the ten *Bgt* isolates.

**Table S6. Gene candidates and annotations in Chinese Spring for regions associated to powdery mildew resistance.** Highlighted genes are the best candidates based on their annotation.

**Table S7. List of all significantly associated SNPs, their position, MAF, effect size beta and p-values for the LandracePLUS panel excluding subsets.**

**Table S8. List of all significantly associated SNPs, their position, MAF, effect size beta and p-values for the subsets of the LandracePLUS panel.**

**Table S9. Raw phenotype data of the LandracePLUS panel infected with ten powdery mildew isolates.** Plates refer to the individual petri dish the replicate was in and Rounds to a batch that was infected at the same time.

**Supplementary file3**

Alleles in the twenty most modern cultivars of the LandracePLUS panel for each potentially novel resistance-associated region. Resistance-associated alleles and accessions with more than 50% of these alleles and a resistance phenotype towards the peak-producing isolate are highlighted in bold.

**Dataset**

**Data S1. Raw genotyping data of the LandracePLUS panel with the TaBW35K SNP array.** Empty cells are labelled “vide”, and wheat accession IDs are extended by “_cell identifier”.

**Data S2. Sample statistics of the raw genotyping data of the LandracePLUS panel.** Total numbers are labelled “nb” and percentage “pc”.

**Data S3. Marker statistics of the raw genotyping data of the LandracePLUS panel.** Total numbers are labelled “nb” and percentage “pc”.

## Notes

### Competing Interest Statement

The authors have declared no competing interest.

### Summary of Updates

We used a curated phenotype input file for all data analyses, where phenotypes with less than three replicates and heterogenous phenotypes were removed. Corresponding analysis and figures have been accordingly updated.

## References

Alemu SK, Huluka AB, Tesfaye K, et al (2021) Genome-wide association mapping identifies yellow rust resistance loci in Ethiopian durum wheat germplasm. PLoS One 16:e0243675. 10.1371/journal.pone.0243675

Alexander DH, Novembre J, Lange K (2009) Fast model-based estimation of ancestry in unrelated individuals. Genome Res 19:1655–1664. 10.1101/gr.094052.109

Athiyannan N, Abrouk M, Boshoff WHP, et al (2022a) Long-read genome sequencing of bread wheat facilitates disease resistance gene cloning. Nat Genet 54:227–231. 10.1038/s41588-022-01022-1

Athiyannan N, Aouini L, Wang Y, Krattinger SG (2022b) Unconventional R proteins in the botanical tribe Triticeae. Essays Biochem 66:561–569. 10.1042/EBC20210081

Aury JM, Engelen S, Istace B, et al (2022) Long-read and chromosome-scale assembly of the hexaploid wheat genome achieves high resolution for research and breeding. Gigascience 11:1–18. 10.1093/gigascience/giac034

Balfourier F, Bouchet S, Robert S, et al (2019) Worldwide phylogeography and history of wheat genetic diversity. Sci Adv 5:. 10.1126/sciadv.aav0536

Bates D, Mächler M, Bolker BM, Walker SC (2015) Fitting linear mixed-effects models using lme4. J Stat Softw 67:1–48. 10.18637/jss.v067.i01

Ben-David R, Xie W, Peleg Z, et al (2010) Identification and mapping of PmG16, a powdery mildew resistance gene derived from wild emmer wheat. Theoretical and Applied Genetics 121:499–510. 10.1007/s00122-010-1326-5

Bhullar NK, Mackay M, Keller B (2010a) Genetic Diversity of the Pm3 Powdery Mildew Resistance Alleles in Wheat Gene Bank Accessions as Assessed by Molecular Markers. Diversity (Basel) 2:768–786. 10.3390/d2050768

Bhullar NK, Street K, Mackay M, et al (2009) Unlocking wheat genetic resources for the molecular identification of previously undescribed functional alleles at the Pm3 resistance locus. Proc Natl Acad Sci U S A 106:9519–9524. 10.1073/pnas.0904152106

Bhullar NK, Zhang Z, Wicker T, Keller B (2010b) Wheat gene bank accessions as a source of new alleles of the powdery mildew resistance gene Pm3: A large scale allele mining project. BMC Plant Biol 10:88. 10.1186/1471-2229-10-88

Blanco A, Gadaleta A, Cenci A, et al (2008) Molecular mapping of the novel powdery mildew resistance gene Pm36 introgressed from Triticum turgidum var. dicoccoides in durum wheat. Theoretical and Applied Genetics 117:135–142. 10.1007/s00122-008-0760-0

Bonjean AP, Angus WJ (2001) The world wheat book: a history of wheat breeding. Lavoisier Publishing Inc.

Bourguet D, Guillemaud T (2016) The Hidden and External Costs of Pesticide Use. Springer, Cham, pp 35–120

Bourras S, Kunz L, Xue M, et al (2019) The AvrPm3-Pm3 effector-NLR interactions control both race-specific resistance and host-specificity of cereal mildews on wheat. Nat Commun 10:1–16. 10.1038/s41467-019-10274-1

Bourras S, McNally KE, Ben-David R, et al (2015) Multiple Avirulence Loci and Allele-Specific Effector Recognition Control the *Pm3* Race-Specific Resistance of Wheat to Powdery Mildew. Plant Cell 27:tpc.15.00171. 10.1105/tpc.15.00171

Bradbury PJ, Zhang Z, Kroon DE, et al (2007) TASSEL: software for association mapping of complex traits in diverse samples. Bioinformatics 23:2633–2635. 10.1093/bioinformatics/btm308

Brown JKM (2015) Durable Resistance of Crops to Disease: A Darwinian Perspective. Annu Rev Phytopathol 53:513–539. 10.1146/annurev-phyto-102313-045914

Browning BL, Zhou Y, Browning SR (2018) A One-Penny Imputed Genome from Next-Generation Reference Panels. Am J Hum Genet 103:338–348. 10.1016/j.ajhg.2018.07.015

Cakir C, Gillespie ME, Scofield SR (2010) Rapid Determination of Gene Function by Virus-induced Gene Silencing in Wheat and Barley. Crop Sci 50:S-77–S-84. 10.2135/cropsci2009.10.0567

CGIAR (2023) Genebank Platform. https://www.genebanks.org/resources/crops/wheat/. Accessed 8 Mar 2023

Chao K, Su W, Wu L, et al (2019) Molecular Mapping of a Recessive Powdery Mildew Resistance Gene in Wheat Cultivar Tian Xuan 45 Using Bulked Segregant Analysis with Polymorphic Single Nucleotide Polymorphism Relative Ratio Distribution. Phytopathology 109:828–838. 10.1094/PHYTO-03-18-0092-R

Chen F, Jia H, Zhang X, et al (2019) Positional cloning of PmCH1357 reveals the origin and allelic variation of the Pm2 gene for powdery mildew resistance in wheat. Crop Journal 7:771–783. 10.1016/j.cj.2019.08.004

Chen PD, Qi LL, Zhou B, et al (1995) Development and molecular cytogenetic analysis of wheat-Haynaldia villosa 6VS/6AL translocation lines specifying resistance to powdery mildew. Theoretical and Applied Genetics 91:1125–1128. 10.1007/BF00223930

Chhuneja P, Yadav B, Stirnweis D, et al (2015) Fine mapping of powdery mildew resistance genes PmTb7A.1 and PmTb7A.2 in Triticum boeoticum (Boiss.) using the shotgun sequence assembly of chromosome 7AL. Theoretical and Applied Genetics 128:2099–2111. 10.1007/s00122-015-2570-5

Danecek P, Auton A, Abecasis G, et al (2011) The variant call format and VCFtools. Bioinformatics 27:2156–2158. 10.1093/bioinformatics/btr330

de Vries A, Ripley BD (2022) Create Dendrograms and Tree Diagrams Using “ggplot2”

Dodds PN, Rathjen JP (2010) Plant immunity: Towards an integrated view of plant-pathogen interactions. Nat Rev Genet 11:539–548. 10.1038/nrg2812

Dong Z, Tian X, Ma C, et al (2020) Physical Mapping of Pm57, a Powdery Mildew Resistance Gene Derived from Aegilops searsii. Int J Mol Sci 21:322. 10.3390/ijms21010322

Dormann CF, Schweiger O, Augenstein I, et al (2007) Effects of landscape structure and land-use intensity on similarity of plant and animal communities. Global Ecology and Biogeography 16:774–787. 10.1111/j.1466-8238.2007.00344.x

Dracatos PM, Lu J, Sánchez-Martín J, Wulff BBH (2023) Resistance that stacks up: engineering rust and mildew disease control in the cereal crops wheat and barley. Plant Biotechnol J 21:1938–1951. 10.1111/pbi.14106

Driscoll CJ, Bielig LM (1968) MAPPING OF THE TRANSEC WHEAT-RYE TRANSLOCATION. Canadian Journal of Genetics and Cytology 10:421–425. 10.1139/g68-056

EU 2020 Farm to Fork. https://ec.europa.eu/commission/presscorner/detail/en/qanda_22_3694. Accessed 8 Mar 2023

Evenson RE, Gollin D (2003) Assessing the impact of the Green Revolution, 1960 to 2000. Science (1979) 300:758–762. 10.1126/science.1078710

FAO (2020) Food and Agriculture Organization of the United Nations, FAOSTAT statistics database, Food Balances. https://www.fao.org/faostat/en/#data/FBS. Accessed 8 Mar 2023

Flor HH (1971) Current status of the gene-for-gene concept. Annu Rev Phytopathol 9:275– 296

Fu B, Chen Y, Li N, et al (2013) pmX: A recessive powdery mildew resistance gene at the Pm4 locus identified in wheat landrace Xiaohongpi. Theoretical and Applied Genetics 126:913–921. 10.1007/s00122-012-2025-1

Galili T (2015) dendextend: an R package for visualizing, adjusting and comparing trees of hierarchical clustering. Bioinformatics 31:3718–3720. 10.1093/bioinformatics/btv428

Gao H, Zhu F, Jiang Y, et al (2012) Genetic analysis and molecular mapping of a new powdery mildew resistant gene Pm46 in common wheat. Theoretical and Applied Genetics 125:967–973. 10.1007/s00122-012-1886-7

Glémin S, Scornavacca C, Dainat J, et al (2019) Pervasive hybridizations in the history of wheat relatives. Sci Adv 5:. 10.1126/sciadv.aav9188

Gloss AD, Vergnol A, Morton TC, et al (2022) Genome-wide association mapping within a local Arabidopsis thaliana population more fully reveals the genetic architecture for defensive metabolite diversity. Philosophical Transactions of the Royal Society B: Biological Sciences 377:. 10.1098/rstb.2020.0512

Govta N, Polda I, Sela H, et al (2022) Genome-Wide Association Study in Bread Wheat Identifies Genomic Regions Associated with Grain Yield and Quality under Contrasting Water Availability. Int J Mol Sci 23:. 10.3390/ijms231810575

Haas M, Schreiber M, Mascher M (2019) Domestication and crop evolution of wheat and barley: Genes, genomics, and future directions. J Integr Plant Biol 61:204–225. 10.1111/jipb.12737

Han J, Zhang L-S, Li G-Q, et al (2009) Molecular Mapping of Powdery Mildew Resistance Gene MlWE18 in Wheat Originated from Wild Emmer (Triticum turgidum var. dicoccoides). ACTA AGRONOMICA SINICA 35:1791–1797. 10.3724/sp.j.1006.2009.01791

Hao M, Liu M, Luo J, et al (2018) Introgression of powdery mildew resistance gene pm56 on rye chromosome arm 6rs into wheat. Front Plant Sci 9:1040. 10.3389/fpls.2018.01040

He H, Ji Y, Zhu S, et al (2017) Genetic, Physical and Comparative Mapping of the Powdery Mildew Resistance Gene Pm21 Originating from Dasypyrum villosum. Front Plant Sci 8:1914. 10.3389/fpls.2017.01914

He H, Zhu S, Zhao R, et al (2018) Pm21, Encoding a Typical CC-NBS-LRR Protein, Confers Broad-Spectrum Resistance to Wheat Powdery Mildew Disease. Mol Plant 11:879– 882. 10.1016/j.molp.2018.03.004

Hewitt T, Müller MC, Molnár I, et al (2021) A highly differentiated region of wheat chromosome 7AL encodes a *Pm1a* immune receptor that recognizes its corresponding *AvrPm1a* effector from *Blumeria graminis*. New Phytologist 229:2812–2826. 10.1111/nph.17075

Hinterberger V, Douchkov D, Lück S, et al (2022) Mining for New Sources of Resistance to Powdery Mildew in Genetic Resources of Winter Wheat. Front Plant Sci 13:366. 10.3389/fpls.2022.836723

Hsam SLK, Huang XQ, Zeller FJ (2001) Chromosomal location of genes for resistance to powdery mildew in common wheat (Triticum aestivum L. em Thell.) 6. Alleles at the Pm5 locus. Theoretical and Applied Genetics 102:127–133. 10.1007/s001220051627

Hu TZ, Li HJ, Liu ZJ, et al (2008) Identification and Molecular Mapping of the Powdery Mildew Resistance Gene in Wheat Cultivar Yumai 66. ACTA AGRONOMICA SINICA 34:545–550. 10.3724/sp.j.1006.2008.00545

Huang XQ, Hsam SLK, Zeller FJ (2002) Chromosomal location of genes for resistance to powdery mildew in Chinese wheat lines Jieyan 94-1-1 and Siyan 94-1-2. Hereditas 136:212–218. 10.1034/j.1601-5223.2002.t01-1-1360306.x

Iwgsc, Appels R, Eversole K, et al (2018) Shifting the limits in wheat research and breeding using a fully annotated reference genome. Science (1979) 361:. 10.1126/science.aar7191

Ji X, Xie C, Ni Z, et al (2008) Identification and genetic mapping of a powdery mildew resistance gene in wild emmer (Triticum dicoccoides) accession IW72 from Israel. Euphytica 159:385–390. 10.1007/s10681-007-9540-1

Jin Y, Xue F, Zhou Y, et al (2020) Fine-Mapping of the Powdery Mildew Resistance Gene mlxbd in the Common Wheat Landrace Xiaobaidong. Plant Dis 104:1231–1238. 10.1094/PDIS-07-19-1347-RE

Kale SM, Schulthess AW, Padmarasu S, et al (2022) A catalogue of resistance gene homologs and a chromosome-scale reference sequence support resistance gene mapping in winter wheat. Plant Biotechnol J 20:1730–1742. 10.1111/pbi.13843

Kaur N, Street K, Mackay M, et al (2008) Molecular approaches for characterization and use of natural disease resistance in wheat. In: European Journal of Plant Pathology. Springer, pp 387–397

Keilwagen J, Lehnert H, Berner T, et al (2022) Detecting major introgressions in wheat and their putative origins using coverage analysis. Sci Rep 12:1908. 10.1038/s41598-022-05865-w

Kim SH, Hong JK, Lee SC, et al (2004) CAZFP1, Cys2/His2-type zinc-finger transcription factor gene functions as a pathogen-induced early-defense gene in Capsicum annuum. Plant Mol Biol 55:883–904. 10.1007/s11103-004-2151-5

Kloppe T, Whetten RB, Kim S, et al (2023) Two pathogen loci determine *Blumeria graminis* f. sp. *tritici* virulence to wheat resistance gene *Pm1a*. New Phytologist. 10.1111/nph.18809

Knaus BJ, Grünwald NJ (2017) VCFR: a package to manipulate and visualize variant call format data in R. In: Molecular Ecology Resources. Blackwell Publishing Ltd, pp 44–53

Koller T, Brunner S, Herren G, et al (2019) Field grown transgenic Pm3e wheat lines show powdery mildew resistance and no fitness costs associated with high transgene expression. Transgenic Res 28:9–20. 10.1007/s11248-018-0099-5

Kolodziej MC, Singla J, Sánchez-Martín J, et al (2021) A membrane-bound ankyrin repeat protein confers race-specific leaf rust disease resistance in wheat. Nat Commun 12:1–12. 10.1038/s41467-020-20777-x

Komáromi J, Jankovics T, Fábián A, et al (2016) Powdery Mildew Resistance in Wheat Cultivar Mv Hombár is Conferred by a New Gene, PmHo. Phytopathology 106:1326– 1334. 10.1094/PHYTO-03-16-0152-R

Kunz L, Sotiropoulos AG, Graf J, et al (2023) The broad use of the Pm8 resistance gene in wheat resulted in hypermutation of the AvrPm8 gene in the powdery mildew pathogen. BMC Biol 21:29. 10.1186/s12915-023-01513-5

Li G, Cowger C, Wang X, et al (2019) Characterization of Pm65, a new powdery mildew resistance gene on chromosome 2AL of a facultative wheat cultivar. Theoretical and Applied Genetics 132:2625–2632. 10.1007/s00122-019-03377-2

Li M, Dong L, Li B, et al (2020a) A CNL protein in wild emmer wheat confers powdery mildew resistance. New Phytologist 228:1027–1037. 10.1111/nph.16761

Li Y, Shi X, Hu J, et al (2020b) Identification of a recessive gene PMQ conferring resistance to powdery mildew in wheat Landrace qingxinmai using BSR-seq analysis. Plant Dis 104:743–751. 10.1094/PDIS-08-19-1745-RE

Lillemo M, Asalf B, Singh RP, et al (2008) The adult plant rust resistance loci Lr34/Yr18 and Lr46/Yr29 are important determinants of partial resistance to powdery mildew in bread wheat line Saar. Theoretical and Applied Genetics 116:1155–1166. 10.1007/s00122-008-0743-1

Liu J, He Z, Rasheed A, et al (2017) Genome-wide association mapping of black point reaction in common wheat (Triticum aestivum L.). BMC Plant Biol 17:220. 10.1186/s12870-017-1167-3

Liu Y, Chen J, Yin C, et al (2023) A high-resolution genotype–phenotype map identifies the TaSPL17 controlling grain number and size in wheat. Genome Biol 24:1–28. 10.1186/s13059-023-03044-2

Loh YT, Martin GB (1995) The Pto bacterial resistance gene and the Fen insecticide sensitivity gene encode functional protein kinases with serine/threonine specificity. Plant Physiol 108:1735–1739. 10.1104/pp.108.4.1735

Lopes MS, El-Basyoni I, Baenziger PS, et al (2015) Exploiting genetic diversity from landraces in wheat breeding for adaptation to climate change. J Exp Bot 66:3477–3486. 10.1093/jxb/erv122

Luo PG, Luo HY, Chang ZJ, et al (2009) Characterization and chromosomal location of Pm40 in common wheat: a new gene for resistance to powdery mildew derived from Elytrigia intermedium. Theoretical and Applied Genetics 118:1059–1064. 10.1007/s00122-009-0962-0

Lutz J, Hsam SLK, Limpert E, Zeller FJ (1995) Chromosomal location of powdery mildew resistance genes in triticum aestivum L. (common wheat). 2. genes pm2 and pm19 from aegi/ops squarrosa L. Heredity (Edinb) 74:152–156. 10.1038/hdy.1995.22

Lutz J, Limpert E, Bartos P, Zeller FJ (1992) Identification of Powdery Mildew Resistance Genes in Common Wheat (Triticum aestivum L.). I. Czechoslovakian Cultivars. Plant Breeding 108:33–39. 10.1111/j.1439-0523.1992.tb00097.x

Mackay M, Street K (2004) Focused identification of germplasm strategy — FIGS. In: Proceedings of the 54th Australian Cereal Chemistry Conference and the 11th Wheat Breeders’ Assembly. Cereal Chemestry Division, Royal Australian Chemical Institute (RACI), Melbourne, Victoria, Australia., pp 138–141

Manser B, Koller T, Praz CR, et al (2021) Identification of specificity-defining amino acids of the wheat immune receptor Pm2 and powdery mildew effector AvrPm2. The Plant Journal 106:993–1007. 10.1111/tpj.15214

Mantel N, Valand RS (1970) A Technique of Nonparametric Multivariate Analysis. Biometrics 26:547. 10.2307/2529108

Marees AT, de Kluiver H, Stringer S, et al (2018) A tutorial on conducting genome-wide association studies: Quality control and statistical analysis. Int J Methods Psychiatr Res 27:. 10.1002/mpr.1608

Marone D, Russo MA, Mores A, et al (2021) Importance of landraces in cereal breeding for stress tolerance. Plants 10:1267. 10.3390/plants10071267

Martin GB, Brommonschenkel SH, Chunwongse J, et al (1993) Map-based cloning of a protein kinase gene conferring disease resistance in tomato. Science (1979) 262:1432–1436. 10.1126/science.7902614

Mascher M, Schreiber M, Scholz U, et al (2019) Genebank genomics bridges the gap between the conservation of crop diversity and plant breeding. Nat Genet 51:1076– 1081. 10.1038/s41588-019-0443-6

Maxwell JJ, Lyerly JH, Srnic G, et al (2010) MlAB10 : A Triticum turgidum Subsp. dicoccoides Derived Powdery Mildew Resistance Gene Identified in Common Wheat. Crop Sci 50:2261–2267. 10.2135/cropsci2010.04.0195

McDonald BA, Linde C (2002) Pathogen population genetics, evolutionary potential, and durable resistance. Annu Rev Phytopathol 40:349–379. 10.1146/annurev.phyto.40.120501.101443

McIntosh RA, Yamazaki Y, Dubcovsky J, et al (2013) Catalogue of Gene Symbols for Wheat. In: Komugi wheat genetic resources database. https://wheat.pw.usda.gov/GG3/wgc. Accessed 8 Mar 2023

Mohler V, Bauer C, Schweizer G, et al (2013) Pm50: A new powdery mildew resistance gene in common wheat derived from cultivated emmer. J Appl Genet 54:259–263. 10.1007/s13353-013-0158-9

Mohler V, Zeller FJ, Wenzel G, Hsam SLK (2005) Chromosomal location of genes for resistance to powdery mildew in common wheat (Triticum aestivum L. em Thell.). 9. Gene MlZec1 from the Triticum dicoccoides-derived wheat line Zecoi-1. Euphytica 142:161–167. 10.1007/s10681-005-1251-x

Mu Y, Gong W, Qie Y, et al (2022) Identification of the powdery mildew resistance gene in wheat breeding line Yannong 99102-06188 via bulked segregant exome capture sequencing. Front Plant Sci 13:3277. 10.3389/fpls.2022.1005627

Müller K, Wickham H (2022) tibble: Simple Data Frames. https://CRAN.R-project.org/package=tibble

Müller MC, Kunz L, Schudel S, et al (2022) Ancient variation of the AvrPm17 gene in powdery mildew limits the effectiveness of the introgressed rye Pm17 resistance gene in wheat. Proc Natl Acad Sci U S A 119:e2108808119. 10.1073/pnas.2108808119

Müller T, Schierscher-Viret B, Fossati D, et al (2018) Unlocking the diversity of genebanks: whole-genome marker analysis of Swiss bread wheat and spelt. Theoretical and Applied Genetics 131:407–416. 10.1007/s00122-017-3010-5

Mundt CC (2014) Durable resistance: A key to sustainable management of pathogens and pests. Infection, Genetics and Evolution 27:446–455. 10.1016/j.meegid.2014.01.011

Mundt CC (2018) Pyramiding for resistance durability: Theory and practice. Phytopathology 108:792–802. 10.1094/PHYTO-12-17-0426-RVW

Myles S, Peiffer J, Brown PJ, et al (2009) Association mapping: Critical considerations shift from genotyping to experimental design. Plant Cell 21:2194–2202. 10.1105/tpc.109.068437

Niu JS, Wang BQ, Wang YH, et al (2008) Chromosome location and microsatellite markers linked to a powdery mildew resistance gene in wheat line Lankao 90(6). Plant Breeding 127:346–349. 10.1111/j.1439-0523.2007.01480.x

Oksanen J, Simpson Gavin L., Blanchet F. G., et al (2022) vegan: Community Ecology Package. https://CRAN.R-project.org/package=vegan. Accessed 25 May 2023

Ouyang S, Zhang D, Han J, et al (2014) Fine Physical and Genetic Mapping of Powdery Mildew Resistance Gene MlIW172 Originating from Wild Emmer (Triticum dicoccoides). PLoS One 9:e100160. 10.1371/journal.pone.0100160

Özkan H, Brandolini A, Schäfer-Pregl R, Salamini F (2002) AFLP Analysis of a Collection of Tetraploid Wheats Indicates the Origin of Emmer and Hard Wheat Domestication in Southeast Turkey. Mol Biol Evol 19:1797–1801. 10.1093/oxfordjournals.molbev.a004002

Paux E, Lafarge S, Balfourier F, et al (2022) Breeding for Economically and Environmentally Sustainable Wheat Varieties: An Integrated Approach from Genomics to Selection. Biology (Basel) 11:149. 10.3390/biology11010149

Pedersen TL (2020) patchwork: The Composer of Plots. https://CRAN.R-project.org/package=patchwork

Perugini LD, Murphy JP, Marshall D, Brown-Guedira G (2008) Pm37, a new broadly effective powdery mildew resistance gene from Triticum timopheevii. Theoretical and Applied Genetics 116:417–425. 10.1007/s00122-007-0679-x

Petersen S, Lyerly JH, Worthington ML, et al (2015) Mapping of powdery mildew resistance gene Pm53 introgressed from Aegilops speltoides into soft red winter wheat. Theoretical and Applied Genetics 128:303–312. 10.1007/s00122-014-2430-8

Peusha H, Enno T, Priilinn O (2004) Chromosomal Location of Powdery Mildew Resistance Genes and Cytogenetic Analysis of Meiosis in Common Wheat Cultivar Meri. Hereditas 132:29–34. 10.1111/j.1601-5223.2000.00029.x

Praz CR, Bourras S, Zeng F, et al (2017) AvrPm2 encodes an RNase-like avirulence effector which is conserved in the two different specialized forms of wheat and rye powdery mildew fungus. New Phytologist 213:1301–1314. 10.1111/nph.14372

Pugsley AT, Carter M V. (1953) The resistance of twelve varieties of triticum vulgare to erysiphe graminis tritici. Aust J Biol Sci 6:335–346. 10.1071/BI9530335

Purcell S, Neale B, Todd-Brown K, et al (2007) PLINK: A tool set for whole-genome association and population-based linkage analyses. Am J Hum Genet 81:559–575. 10.1086/519795

Qie Y, Sheng Y, Xu H, et al (2019) Identification of a new powdery mildew resistance gene pmDHT at or closely linked to the Pm5 locus in the Chinese wheat landrace dahongtou. Plant Dis 103:2645–2651. 10.1094/PDIS-02-19-0401-RE

Qiu YC, Zhou RH, Kong XY, et al (2005) Microsatellite mapping of a Triticum urartu Tum. derived powdery mildew resistance gene transferred to common wheat (Triticum aestivum L.). Theoretical and Applied Genetics 111:1524–1531. 10.1007/s00122-005-0081-5

R Core Team (2022) R: A language and environment for statistical computing. https://www.r-project.org/

Reif JC, Zhang P, Dreisigacker S, et al (2005) Wheat genetic diversity trends during domestication and breeding. Theoretical and Applied Genetics 110:859–864. 10.1007/s00122-004-1881-8

Rimbert H, Darrier B, Navarro J, et al (2018) High throughput SNP discovery and genotyping in hexaploid wheat. PLoS One 13:. 10.1371/journal.pone.0186329

Robinson JT, Thorvaldsdóttir H, Winckler W, et al (2011) Integrative genomics viewer. Nat Biotechnol 29:24–26. 10.1038/nbt.1754

Sánchez-Martín J, Keller B (2021) NLR immune receptors and diverse types of non-NLR proteins control race-specific resistance in Triticeae. Curr Opin Plant Biol 62:102053. 10.1016/j.pbi.2021.102053

Sánchez-Martín J, Steuernagel B, Ghosh S, et al (2016) Rapid gene isolation in barley and wheat by mutant chromosome sequencing. Genome Biol 17:1–7. 10.1186/s13059-016-1082-1

Sánchez-Martín J, Widrig V, Herren G, et al (2021) Wheat Pm4 resistance to powdery mildew is controlled by alternative splice variants encoding chimeric proteins. Nat Plants 1–15. 10.1038/s41477-021-00869-2

Sansaloni C, Franco J, Santos B, et al (2020) Diversity analysis of 80,000 wheat accessions reveals consequences and opportunities of selection footprints. Nat Commun 11:4572. 10.1038/s41467-020-18404-w

Sato K, Abe F, Mascher M, et al (2021) Chromosome-scale genome assembly of the transformation-amenable common wheat cultivar ’Fielder. DNA Research 28:. 10.1093/dnares/dsab008

Savary S, Willocquet L, Pethybridge SJ, et al (2019) The global burden of pathogens and pests on major food crops. Nat Ecol Evol 3:430–439. 10.1038/s41559-018-0793-y

Schulthess AW, Kale SM, Liu F, et al (2022) Genomics-informed prebreeding unlocks the diversity in genebanks for wheat improvement. Nat Genet 54:1544–1552. 10.1038/s41588-022-01189-7

Sharma R, Mahanty B, Mishra R, Joshi RK (2021) Genome wide identification and expression analysis of pepper C2H2 zinc finger transcription factors in response to anthracnose pathogen Colletotrichum truncatum. 3 Biotech 11:118. 10.1007/s13205-020-02601-x

Shi AN, Leath S, Murphy JP (1998) A major gene for powdery mildew resistance transferred to common wheat from wild einkorn wheat. Phytopathology 88:144–147. 10.1094/PHYTO.1998.88.2.144

Singh BK, Delgado-Baquerizo M, Egidi E, et al (2023) Climate change impacts on plant pathogens, food security and paths forward. Nat Rev Microbiol 21:640–656

Singh SP, Hurni S, Ruinelli M, et al (2018) Evolutionary divergence of the rye Pm17 and Pm8 resistance genes reveals ancient diversity. Plant Mol Biol 98:249–260. 10.1007/s11103-018-0780-3

Singrün C, Hsam SLK, Zeller FJ, et al (2004) Localization of a novel recessive powdery mildew resistance gene from common wheat line RD30 in the terminal region of chromosome 7AL. Theoretical and Applied Genetics 109:210–214. 10.1007/s00122-004-1619-7

Sotiropoulos AG, Arango-Isaza E, Ban T, et al (2022) Global genomic analyses of wheat powdery mildew reveal association of pathogen spread with historical human migration and trade. Nat Commun 13:16. 10.1038/s41467-022-31975-0

Stein N, Herren G, Keller B (2001) A new DNA extraction method for high-throughput marker analysis in a large-genome species such as Triticum aestivum. Plant Breeding 120:354–356. 10.1046/j.1439-0523.2001.00615.x

Sun M, Liu Q, Han Y, et al (2022) PmSN15218: A Potential New Powdery Mildew Resistance Gene on Wheat Chromosome 2AL. Front Plant Sci 13:. 10.3389/fpls.2022.931778

Tan C, Li G, Cowger C, et al (2019) Characterization of Pm63, a powdery mildew resistance gene in Iranian landrace PI 628024. Theoretical and Applied Genetics 132:1137–1144. 10.1007/s00122-018-3265-5

Tan C, Li G, Cowger C, et al (2018) Characterization of Pm59, a novel powdery mildew resistance gene in Afghanistan wheat landrace PI 181356. Theoretical and Applied Genetics 131:1145–1152. 10.1007/s00122-018-3067-9

Tanksley SD, McCouch SR (1997) Seed banks and molecular maps: Unlocking genetic potential from the wild. Science (1979) 277:1063–1066. 10.1126/science.277.5329.1063

Tettelin H, Masignani V, Cieslewicz MJ, et al (2005) Genome analysis of multiple pathogenic isolates of Streptococcus agalactiae: Implications for the microbial “pan-genome.” Proc Natl Acad Sci U S A 102:13950–13955. 10.1073/pnas.0506758102

Tian Z-D, Zhang Y, Liu J, Xie C-H (2010) Novel potato C2H2-type zinc finger protein gene, StZFP1, which responds to biotic and abiotic stress, plays a role in salt tolerance. Plant Biol 12:689–697. 10.1111/j.1438-8677.2009.00276.x

Tosa Y, Tsujimoto H, Ogura H (1987) A gene involved in the resistance of wheat to wheatgrass powdery mildew fungus. Genome 29:850–852. 10.1139/g87-145

Uehara Y, Takahashi Y, Berberich T, et al (2005) Tobacco ZFT1, a transcriptional repressor with a Cys2/His 2 type zinc finger motif that functions in spermine-signaling pathway. Plant Mol Biol 59:435–448. 10.1007/s11103-005-0272-0

Uffelmann E, Huang QQ, Munung NS, et al (2021) Genome-wide association studies. Nature Reviews Methods Primers 1:1–21. 10.1038/s43586-021-00056-9

Van Den Burg HA, Tsitsigiannis DI, Rowland O, et al (2008) The F-box protein ACRE189/ACIF1 regulates cell death and defense responses activated during pathogen recognition in tobacco and tomato. Plant Cell 20:697–719. 10.1105/tpc.107.056978

Villa TCC, Maxted N, Scholten M, Ford-Lloyd B (2005) Defining and identifying crop landraces. Plant Genetic Resources 3:373–384. 10.1079/pgr200591

Vleeshouwers VGAA, Oliver RP (2014) Effectors as tools in disease resistance breeding against biotrophic, hemibiotrophic, and necrotrophic plant pathogens. Molecular Plant-Microbe Interactions 27:196–206. 10.1094/MPMI-10-13-0313-IA

Walkowiak S, Gao L, Monat C, et al (2020) Multiple wheat genomes reveal global variation in modern breeding. Nature 588:277–283. 10.1038/s41586-020-2961-x

Wan W, Xiao J, Li M, et al (2020) Fine mapping of wheat powdery mildew resistance gene Pm6 using 2B/2G homoeologous recombinants induced by the ph1b mutant. Theoretical and Applied Genetics 133:1265–1275. 10.1007/s00122-20-03546-8

Wang J, Li Y, Xu F, et al (2022) Candidate powdery mildew resistance gene in wheat landrace cultivar Hongyoumai discovered using SLAF and BSR-seq. BMC Plant Biol 22:83. 10.1186/s12870-022-03448-5

Wang J, Zhang Z (2021) GAPIT Version 3: Boosting Power and Accuracy for Genomic Association and Prediction. Genomics Proteomics Bioinformatics 19:629–640. 10.1016/j.gpb.2021.08.005

Wang W, He H, Gao H, et al (2021) Characterization of the Powdery Mildew Resistance Gene in Wheat Breeding Line KN0816 and Its Evaluation in Marker-Assisted Selection. Plant Dis 105:4042–4050. 10.1094/PDIS-05-21-0896-RE

Wei T, Simko V (2021) R package “corrplot”: Visualization of a Correlation Matrix. https://github.com/taiyun/corrplot

Weigel RR, Pfitzner UM, Gatz C (2005) Interaction of NIMIN1 with NPR1 modulates PR gene expression in Arabidopsis. Plant Cell 17:1279–1291. 10.1105/tpc.104.027441

Wenger AM, Peluso P, Rowell WJ, et al (2019) Accurate circular consensus long-read sequencing improves variant detection and assembly of a human genome. Nat Biotechnol 37:1155–1162. 10.1038/s41587-019-0217-9

Wickham H (2016) ggplot2: Elegant Graphics for Data Analysis. Springer-Verlag New York Wickham H, François R, Henry L, et al (2023) dplyr: A Grammar of Data Manipulation. https://CRAN.R-project.org/paclage=dplyr

Worthington M, Lyerly J, Petersen S, et al (2014) *MlUM15* : an *Aegilops neglecta* -derived Powdery Mildew Resistance Gene in Common Wheat. Crop Sci 54:1397–1406. 10.2135/cropsci2013.09.0634

Wu L, Zhu T, He H, et al (2022) Genetic dissection of the powdery mildew resistance in wheat breeding line LS5082 using BSR-Seq. Crop Journal 10:1120–1130. 10.1016/j.cj.2021.12.008

Wu P, Hu J, Zou J, et al (2019) Fine mapping of the wheat powdery mildew resistance gene Pm52 using comparative genomics analysis and the Chinese Spring reference genomic sequence. Theoretical and Applied Genetics 132:1451–1461. 10.1007/s00122-019-03291-7

Xiao M, Song F, Jiao J, et al (2013) Identification of the gene Pm47 on chromosome 7BS conferring resistance to powdery mildew in the Chinese wheat landrace Hongyanglazi. Theoretical and Applied Genetics 126:1397–1403. 10.1007/s00122-013-2060-6

Xie JZ, Wang L li, Wang Y, et al (2017) Fine mapping of powdery mildew resistance gene PmTm4 in wheat using comparative genomics. J Integr Agric 16:540–550. 10.1016/S2095-3119(16)61377-1

Xing L, Hu P, Liu J, et al (2018) Pm21 from Haynaldia villosa Encodes a CC-NBS-LRR Protein Conferring Powdery Mildew Resistance in Wheat. Mol Plant 11:874–878. 10.1016/j.molp.2018.02.013

Xu H, Yao G, Xiong L, et al (2008) Identification and mapping of pm2026: A recessive powdery mildew resistance gene in an einkorn (Triticum monococcum L.) accession. Theoretical and Applied Genetics 117:471–477. 10.1007/s00122-008-0791-6

Xu W, Li C, Hu L, et al (2011) Identification and molecular mapping of PmHNK54: a novel powdery mildew resistance gene in common wheat. Plant Breeding 130:603–607. 10.1111/j.1439-0523.2011.01882.x

Xu WG, Li CX, Hu L, et al (2010) Molecular mapping of powdery mildew resistance gene PmHNK in winter wheat (Triticum aestivum L.) cultivar Zhoumai 22. Molecular Breeding 26:31–38. 10.1007/s11032-009-9374-8

Xu X, Li Q, Ma Z, et al (2018) Molecular mapping of powdery mildew resistance gene PmSGD in Chinese wheat landrace Shangeda using RNA-seq with bulk segregant analysis. Molecular Breeding 38:1–12. 10.1007/s11032-018-0783-4

Xu X, Liu W, Liu Z, et al (2020) Mapping Powdery Mildew Resistance Gene pmYBL on Chromosome 7B of Chinese Wheat (Triticum aestivum L.) Landrace Youbailan. Plant Dis 104:2411–2417. 10.1094/PDIS-01-20-0118-RE

Xue F, Ji W, Wang C, et al (2012) High-density mapping and marker development for the powdery mildew resistance gene PmAS846 derived from wild emmer wheat (Triticum turgidum var. dicoccoides). Theoretical and Applied Genetics 124:1549–1560. 10.1007/s00122-012-1809-7

Yahiaoui N, Srichumpa P, Dudler R, Keller B (2004) Genome analysis at different ploidy levels allows cloning of the powdery mildew resistance gene Pm3b from hexaploid wheat. The Plant Journal 37:528–538. 10.1046/j.1365-313X.2003.01977.x

Yao G, Zhang J, Yang L, et al (2007) Genetic mapping of two powdery mildew resistance genes in einkorn (Triticum monococcum L.) accessions. Theoretical and Applied Genetics 114:351–358. 10.1007/s00122-006-0438-4

Yin J, Wang L, Zhao J, et al (2020) Genome-wide characterization of the C2H2 zinc-finger genes in Cucumis sativus and functional analyses of four CsZFPs in response to stresses. BMC Plant Biol 20:359. 10.1186/s12870-020-02575-1

Yin JL, Fang ZW, Sun C, et al (2018) Rapid identification of a stripe rust resistant gene in a space-induced wheat mutant using specific locus amplified fragment (SLAF) sequencing. Sci Rep 8:1–9. 10.1038/s41598-018-21489-5

Yu X, Ren S, Zhao L, et al (2018) Molecular mapping of a novel wheat powdery mildew resistance gene Ml92145E8-9 and its application in wheat breeding by marker-assisted selection. Crop Journal 6:621–627. 10.1016/j.cj.2018.04.004

Zeller FJ, Kong L, Hartl L, et al (2002) Chromosomal location of genes for resistance to powdery mildew in common wheat (Triticum aestivum L. em Thell.) 7. Gene Pm29 in line Pova. Euphytica 123:187–194. 10.1023/A:1014944619304

Zeven AC (1998) Landraces: A review of definitions and classifications. Euphytica 104:127–139. 10.1023/A:1018683119237

Zhan H, Li G, Zhang X, et al (2014) Chromosomal Location and Comparative Genomics Analysis of Powdery Mildew Resistance Gene Pm51 in a Putative Wheat-Thinopyrum ponticum Introgression Line. PLoS One 9:e113455. 10.1371/journal.pone.0113455

Zhang D, Zhu K, Dong L, et al (2019) Wheat powdery mildew resistance gene Pm64 derived from wild emmer (Triticum turgidum var. dicoccoides) is tightly linked in repulsion with stripe rust resistance gene Yr5. Crop Journal 7:761–770. 10.1016/j.cj.2019.03.003

Zhang H, Guan H, Li J, et al (2010) Genetic and comparative genomics mapping reveals that a powdery mildew resistance gene Ml3D232 originating from wild emmer co-segregates with an NBS-LRR analog in common wheat (Triticum aestivum L.). Theoretical and Applied Genetics 121:1613–1621. 10.1007/s00122-010-1414-6

Zhang R, Fan Y, Kong L, et al (2018) Pm62, an adult-plant powdery mildew resistance gene introgressed from Dasypyrum villosum chromosome arm 2VL into wheat. Theoretical and Applied Genetics 131:2613–2620. 10.1007/s00122-018-3176-5

Zhang R, Sun B, Chen J, et al (2016) Pm55, a developmental-stage and tissue-specific powdery mildew resistance gene introgressed from Dasypyrum villosum into common wheat. Theoretical and Applied Genetics 129:1975–1984. 10.1007/s00122-016-2753-8

Zhang W, Yu Z, Wang D, et al (2023) Characterization and identification of the powdery mildew resistance gene in wheat breeding line ShiCG15–009. BMC Plant Biol 23:113. 10.1186/s12870-023-04132-y

Zheng X, Levine D, Shen J, et al (2012) A high-performance computing toolset for relatedness and principal component analysis of SNP data. Bioinformatics 28:3326– 3328. 10.1093/bioinformatics/bts606

Zhou X, Stephens M (2012) Genome-wide efficient mixed-model analysis for association studies. Nat Genet 44:821–824. 10.1038/ng.2310

Zhu Z, Zhou R, Kong X, et al (2005) Microsatellite markers linked to 2 powdery mildew resistance genes introgressed from Triticum carthlicum accession PS5 into common wheat. Genome 48:585–590. 10.1139/G05-016

Zhu Z, Zhou R, Kong X, et al (2006) Microsatellite marker identification of a Triticum aestivum-Aegilops umbellulata substitution line with powdery mildew resistance. Euphytica 150:149–153. 10.1007/s10681-006-9103-x

Zou S, Wang H, Li Y, et al (2018) The NB-LRR gene Pm60 confers powdery mildew resistance in wheat. New Phytologist 218:298–309. 10.1111/nph.14964

