## Supplementary_file1 for "A diverse panel of 755 bread wheat accessions harbors untapped genetic diversity in landraces and reveals novel genetic regions conferring powdery mildew resistance"

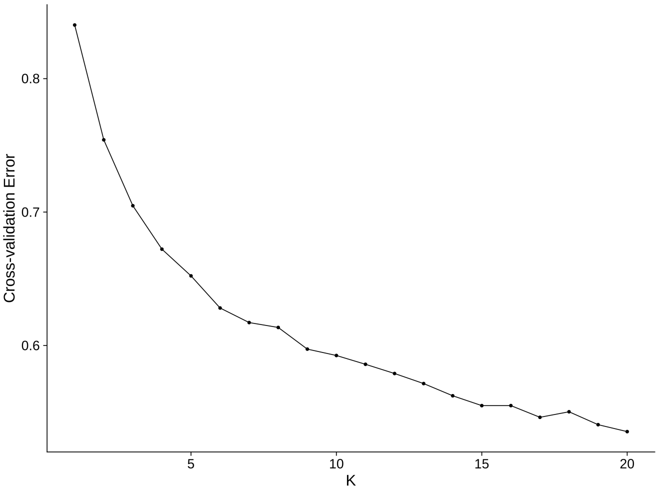


**Figure S1. Cross-validation error of Admixture kinship analysis for K=2 to K=20 for 29,965 SNPs.**

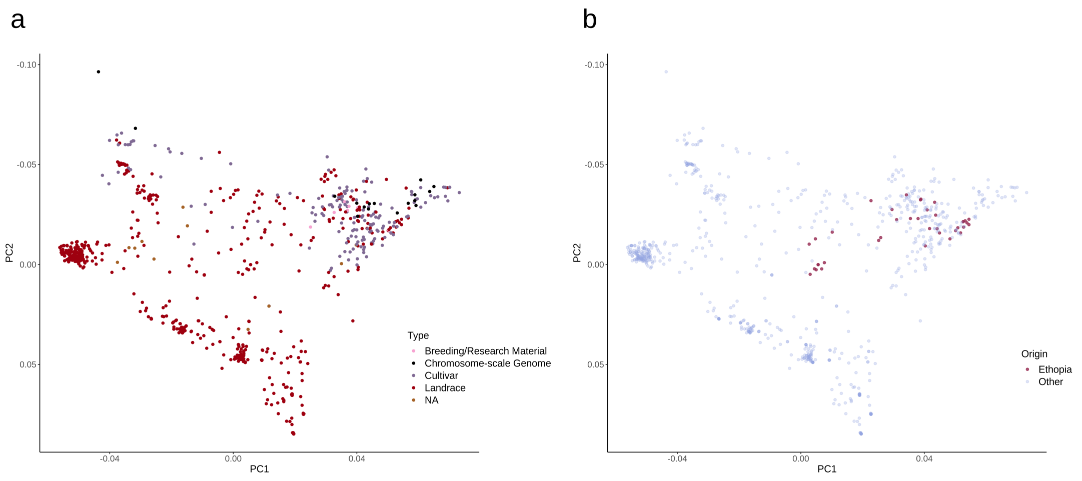


**Figure S2. PCAs from 27,337 SNPs of the LandracePLUS panel including high-quality sequenced genomes with PC1 = 8.4% and PC2 = 5.1%**
**a** Types of wheat accessions are highlighted in different colors. **b** Ethiopian landraces are highlighted by color in comparison to accessions of another origin

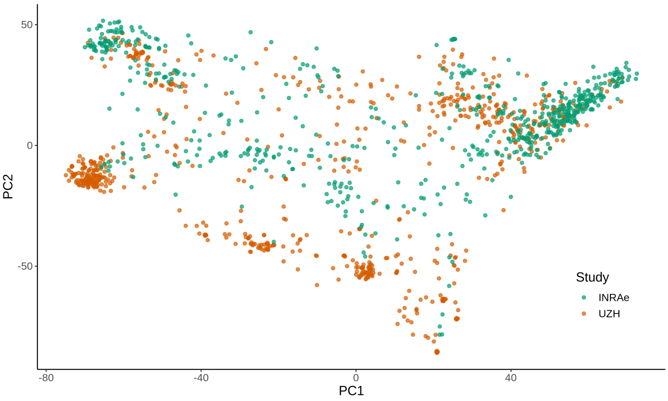


**Figure S3. Genetic diversity of the LandracePLUS panel compared to 632 landraces from an INRAe study that represent the original pool of worldwide hexaploid diversity (Balfourier et al. 2019).**
PCA from the filtered 29,965 polymorphic SNPs that were present in SNP arrays of both studies. PC1 = 8.9%, PC2 = 4.0%. Accessions from the LandracePLUS panel are shown in orange, while INRAe landraces are colored in green.

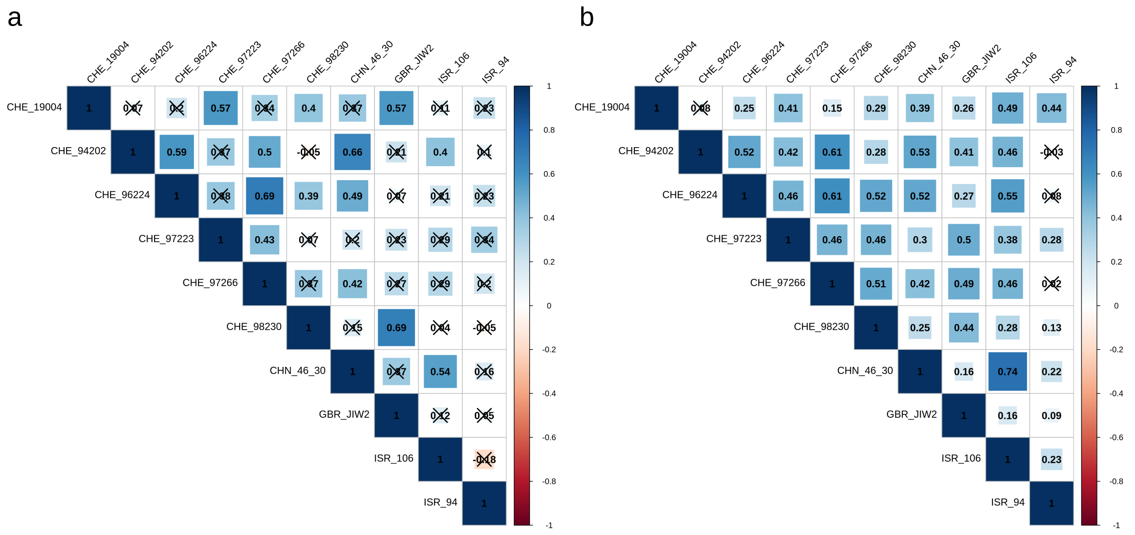

**Figure S4. Correlation matrix showing Pearson correlation of virulence between the ten isolates used for phenotyping.**
**a** Correlation coefficients based on phenotypic variation of the differential lines. **b** Correlation coefficients based on phenotypic variation in the LandracePLUS panel. Correlation coefficients with corresponding p-values below the significance 0.05 are crossed out.

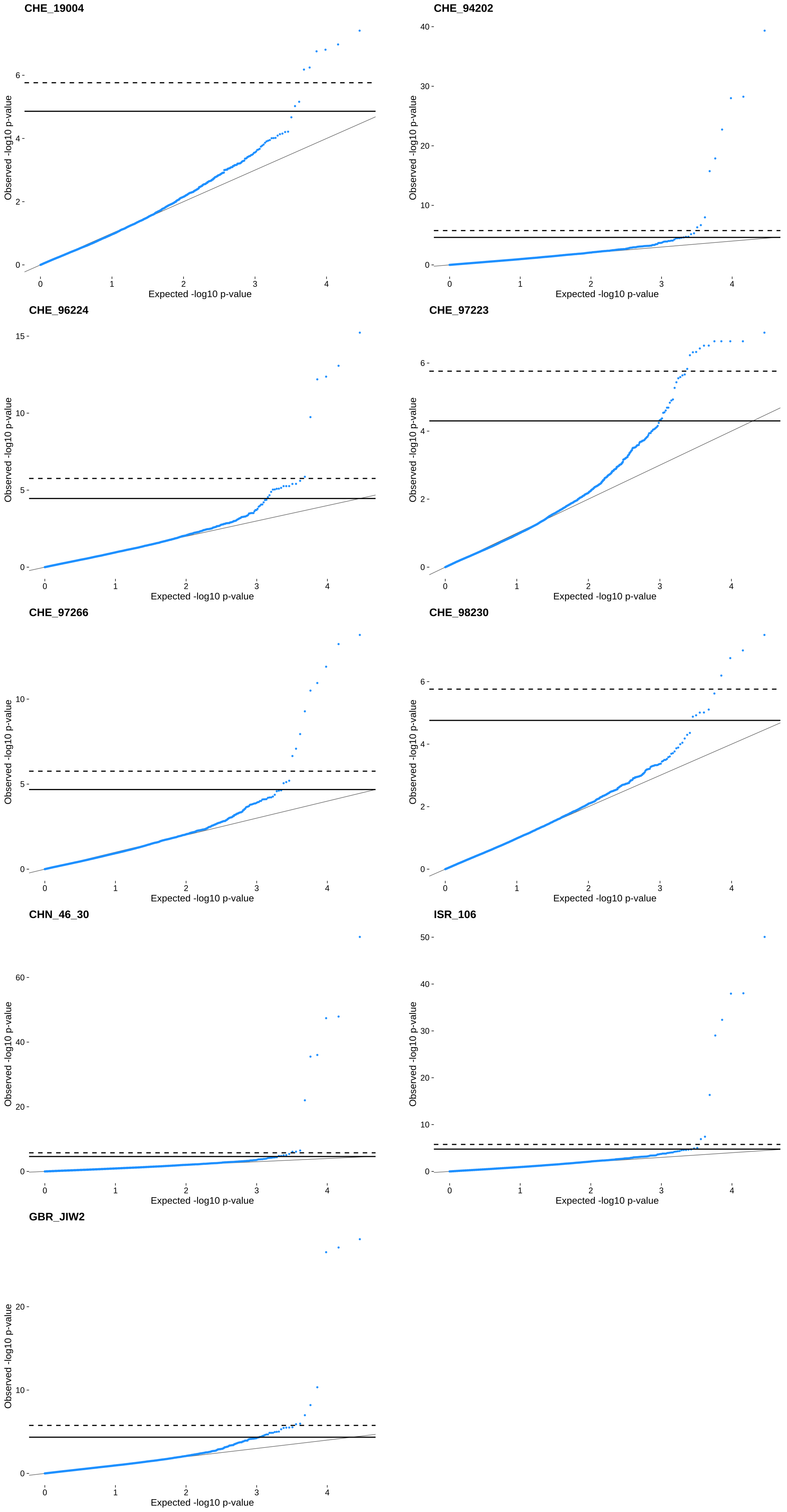


**Figure S5. QQ Plots for GWAS with nine powdery mildew isolates and the LandracePLUS panel.**
Isolate names are depicted in the top-left corner. Solid lines represent the threshold for False discovery rate (FDR) and dashed lines for Bonferroni correction.

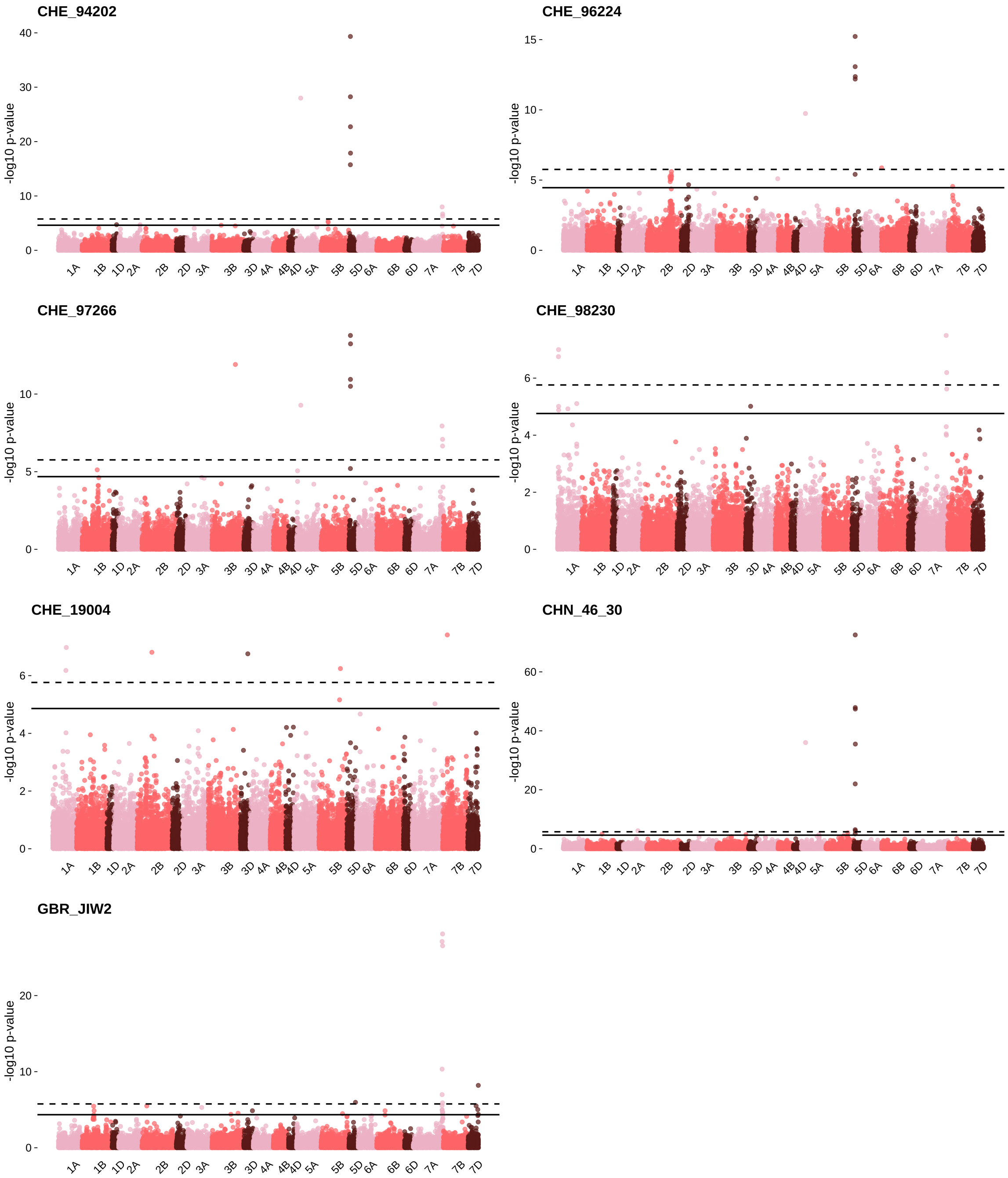


**Figure S6. Manhattan Plots for GWAS with the seven powdery mildew isolates and the LandracePLUS panel that showed a good fit in the QQ plots for the univariate linear mixed model.**
Isolate names are depicted in the top-left corner. Solid lines represent the threshold for False discovery rate (FDR) and dashed lines for Bonferroni correction.

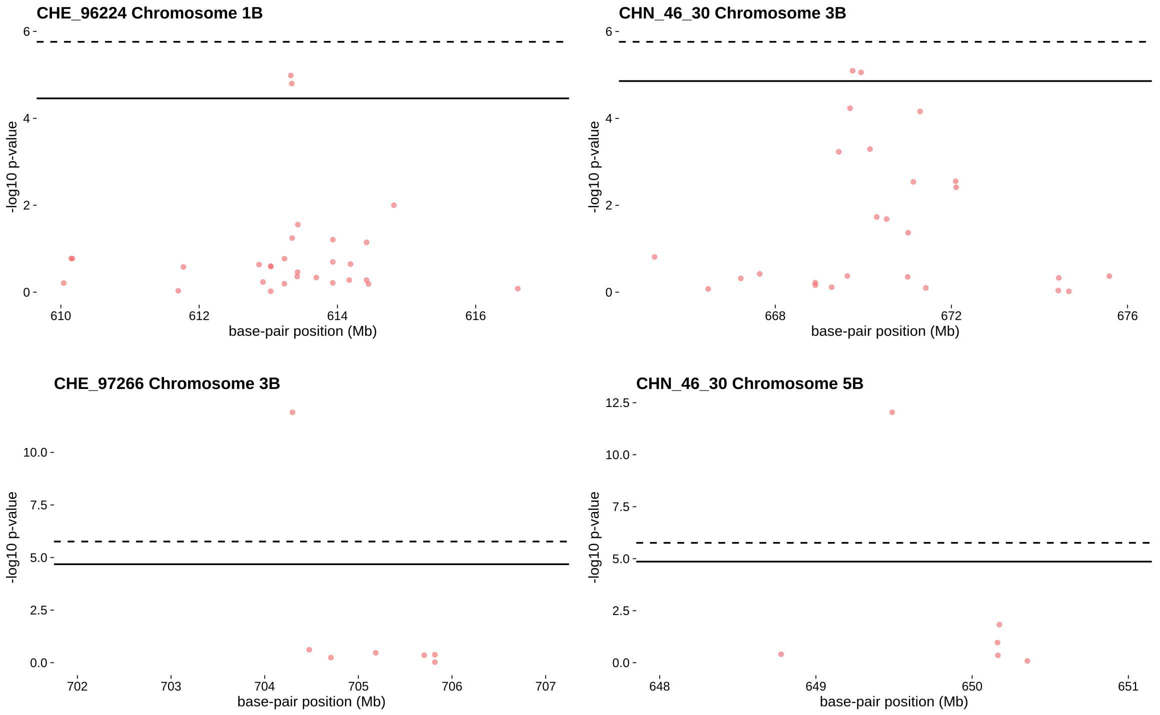


**Figure S7. Manhattan Plots for GWAS of the LandracePLUS panel using a *Pm2* covariate and the isolates that contain *AvrPm2*.** Isolate and chromosomes are depicted in the top-left corner. Solid lines represent the threshold for False discovery rate (FDR) and dashed lines for Bonferroni correction.

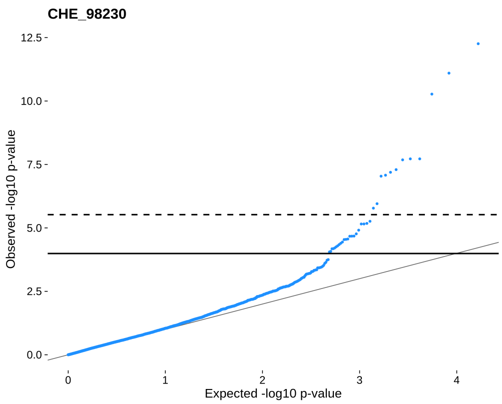


**Figure S8. QQ Plot for GWAS with powdery mildew isolate CHE_98230 and landraces from Pakistan and Iran.**
Isolate names are depicted in the top-left corner. Solid lines represent the threshold for False discovery rate (FDR) and dashed lines for Bonferroni correction.

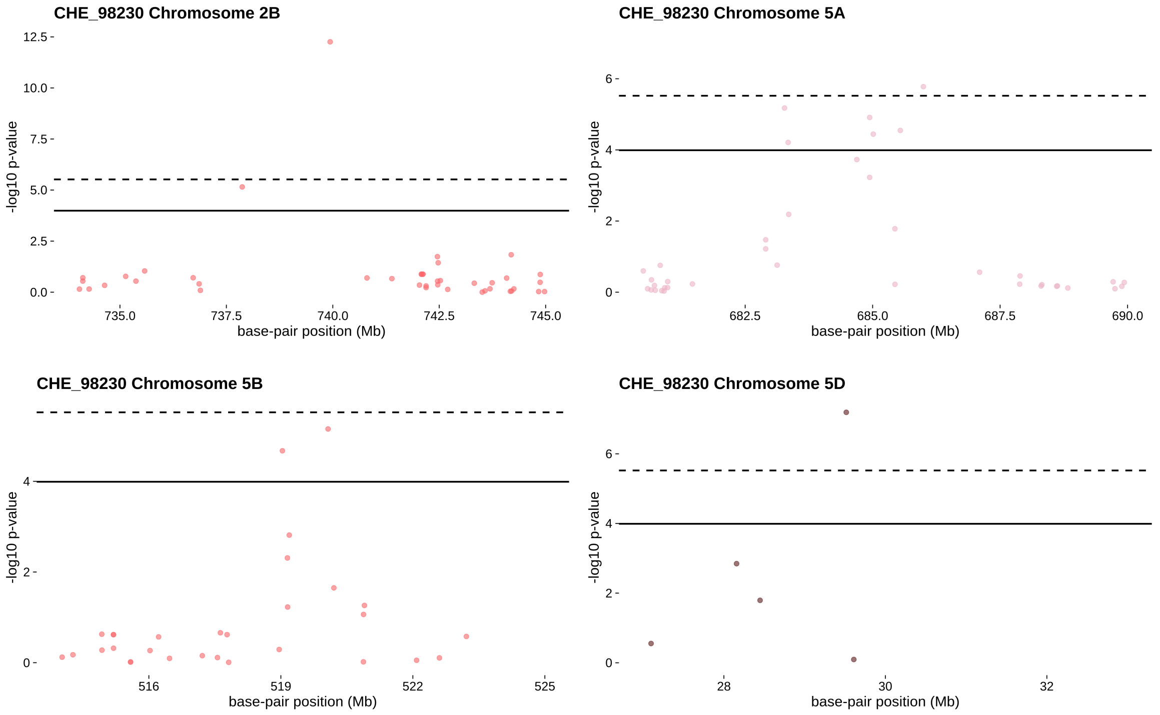


**Figure S9. Manhattan Plots of CHE_98230 resistance-associated loci of accessions from Pakistan and Iran.** Isolate and chromosomes are depicted in the top-left corner. Solid lines represent the threshold for False discovery rate (FDR) and dashed lines for Bonferroni correction.

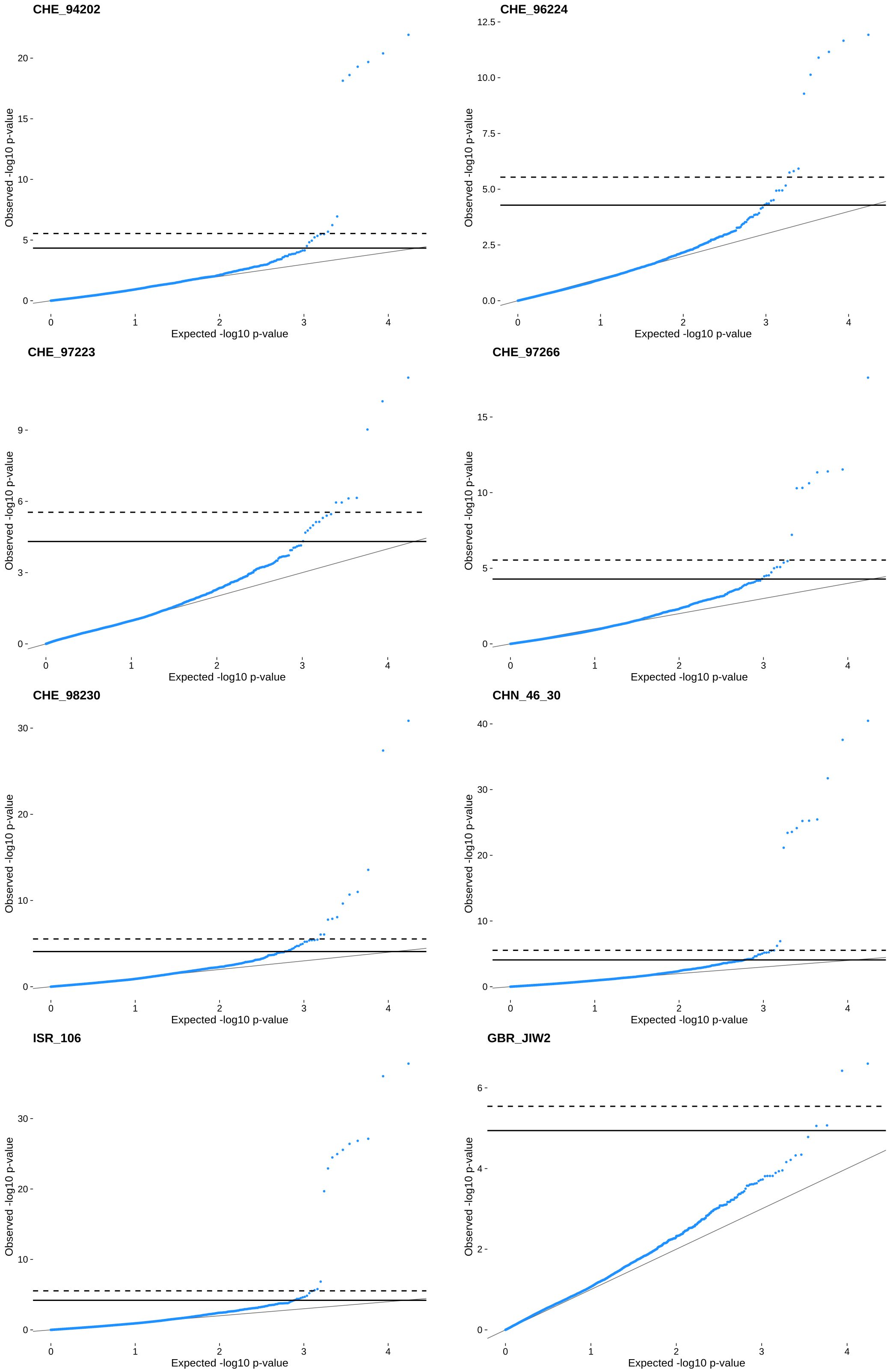


**Figure S10. QQ Plots for GWAS with eight powdery mildew isolates and landraces from Turkey.**
Isolate names are depicted in the top-left corner. Solid lines represent the threshold for False discovery rate (FDR) and dashed lines for Bonferroni correction.

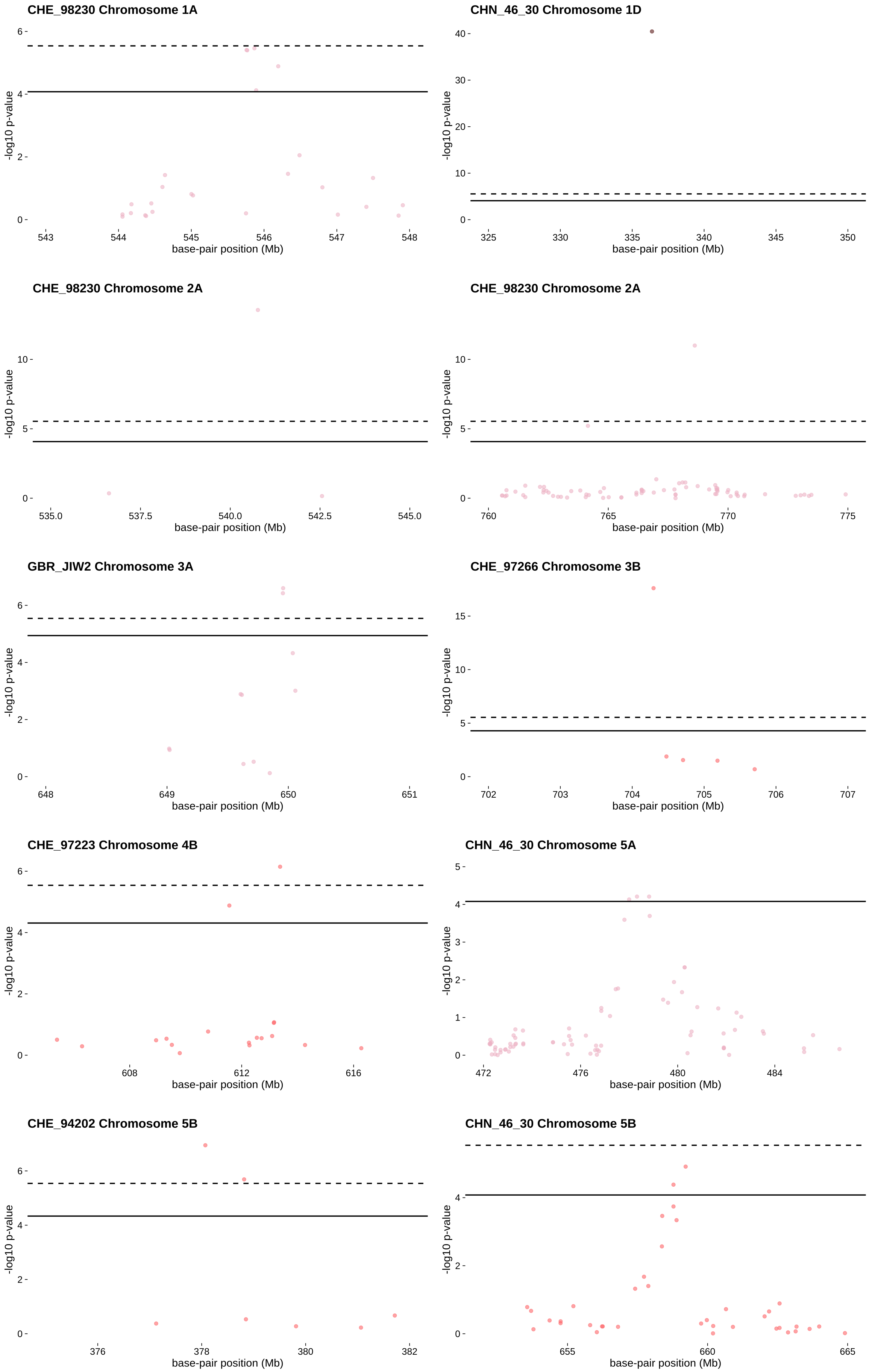

CHE_98230 Chromosome 1A
-log10 p-value
0
2
4
6
543
544
545
546
547
548
base-pair position (Mb)
CHN_46_30 Chromosome 1D
-log10 p-value
0
10
20
30
40
325
330
335
340
345
350
base-pair position (Mb)
CHE_98230 Chromosome 2A
-log10 p-value
0
5
10
535.0
537.5
540.0
542.5
545.0
base-pair position (Mb)
CHE_98230 Chromosome 2A
-log10 p-value
0
5
10
760
765
770
775
base-pair position (Mb)
GBR_JIW2 Chromosome 3A
-log10 p-value
0
2
4
6
648
649
650
651
base-pair position (Mb)
CHE_97266 Chromosome 3B
-log10 p-value
0
5
10
15
702
703
704
705
706
707
base-pair position (Mb)
CHE_97223 Chromosome 4B
-log10 p-value
0
2
4
6
608
612
616
base-pair position (Mb)
CHN_46_30 Chromosome 5A
-log10 p-value
0
1
2
3
4
5
472
476
480
484
base-pair position (Mb)
CHE_94202 Chromosome 5B
-log10 p-value
0
2
4
6
376
378
380
382
base-pair position (Mb)
CHN_46_30 Chromosome 5B
-log10 p-value
0
2
4
655
660
665
base-pair position (Mb)

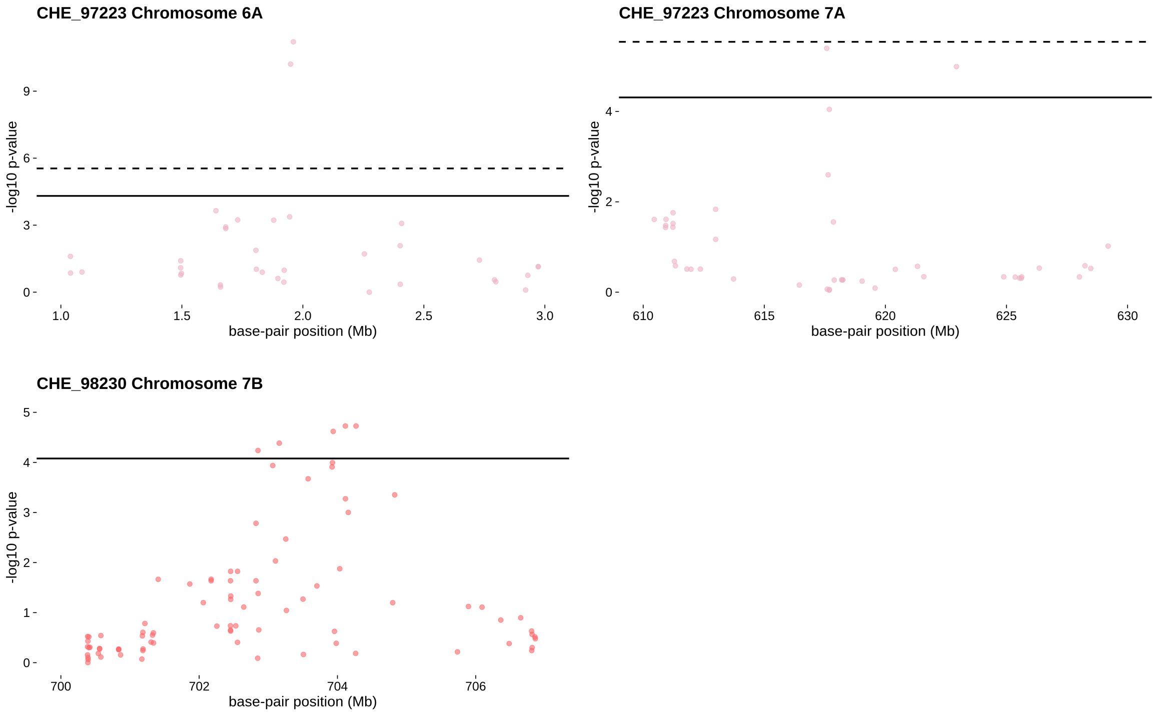


**Figure S11. Manhattan Plots of resistance-associated loci of Turkish accessions.** Isolates and chromosomes are depicted in the top-left corner. Solid lines represent the threshold for False discovery rate (FDR) and dashed lines for Bonferroni correction. Associations that occurred for several isolates are represented by one isolate only.

### References


Balfourier F et al. (2019) Worldwide phylogeography and history of wheat genetic diversity. *Sci Adv* 5.
